# Adaptive capacity of a DNA polymerase clamp-loader ATPase complex

**DOI:** 10.1101/2023.10.18.562948

**Authors:** Subu Subramanian, Weilin Zhang, Siddharth Nimkar, Mazzin Kamel, Michael O’Donnell, John Kuriyan

## Abstract

The ability of mutations to facilitate adaptation is central to evolution. To understand how mutations can lead to functional adaptation in a complex molecular machine, we created a defective version of the T4 clamp-loader complex, which is essential for DNA replication. This variant, which is ∼5000-fold less active than the wildtype, was made by replacing the catalytic domains with those from another phage. A directed-evolution experiment revealed that multiple substitutions to a single negatively-charged residue in the chimeric clamp loader – Asp 86 – restore fitness to within ∼20-fold of wildtype. These mutations remove an adventitious electrostatic repulsive interaction between Asp 86 and the sliding clamp. Deep mutagenesis shows that the reduced fitness of the chimeric clamp loader is compensated for by lysine and arginine substitutions of several DNA-proximal residues in the clamp loader or the sliding clamp. Thus, the fitness decrease of the chimeric clamp loader is caused by a reduction in affinity between the clamp loader and the clamp. Our results demonstrate that there is a latent capacity for increasing affinity of the clamp loader for DNA and the sliding clamp, such that even single point mutations can readily compensate for the loss of function due to suboptimal interactions elsewhere.

---

The evolutionary divergence of proteins is dependent on the ability of mutations to modulate protein function to better align with changing conditions. Protein machines have a remarkable capacity to accept mutations without complete loss of function, and to optimize fitness under changing external constraints. For machines that are critical to the most central processes of life, an important question is how this adaptation is accomplished without extinguishing the capacity of the system to continue living. In this study, we investigate the adaptive capacity of one such critically important protein machine, the DNA polymerase clamp-loader complex.

During genome replication, clamp-loader complexes load ring-shaped sliding clamps onto primer-templated DNA (Kelch et al. 2012). Sliding clamps enable highly processive DNA replication by anchoring the DNA polymerases onto DNA. The clamp loader cycle, illustrated in Fig. 1A, involves three steps: ATP-bound clamp loader binding and opening of the sliding clamp, recognition and binding of the clamp-loader/clamp complex on DNA, and cooperative ATP hydrolysis, leading to the release of the closed clamp around DNA (Kelch 2016).

**Figure 1.**
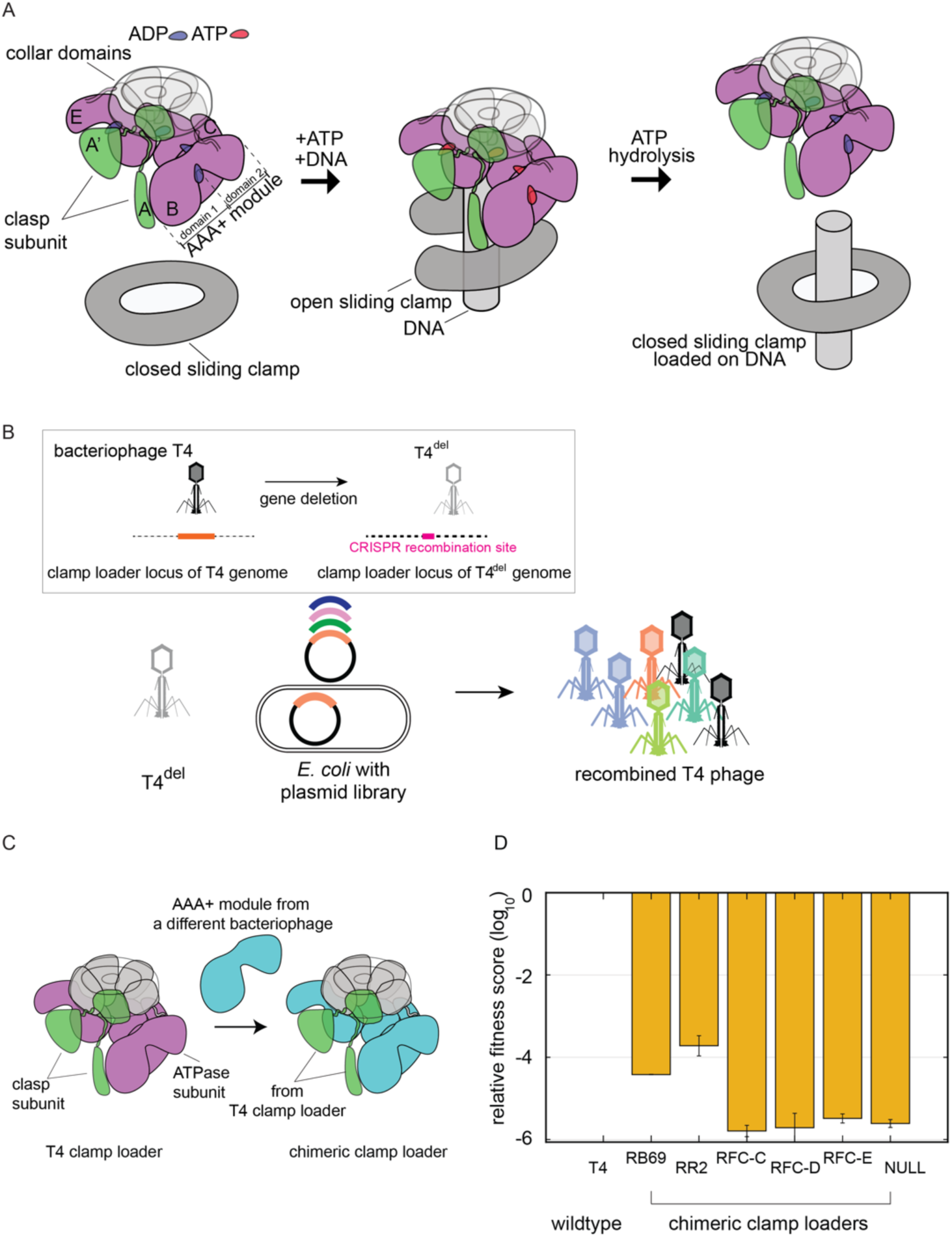
Phage-propagation assay to measure the fitness of clamp-loader variants in T4 bacteriophage. (A) Schematic diagram of the clamp-loading cycle, from left to right, showing key steps of loading the sliding clamp around primer-templated DNA. (B) Schematic of the phage-propagation assay in which T4^del^ infects host cells each of which carry a plasmid-borne copy of variant genes of the T4 clamp loader. Upon infection, the variant genes recombine into the genome of T4^del^, and this genome is replicated using the proteins encoded by the variant genes. The number of copies of the recombinant genomes (and the recombinant phage) is proportional to the activity of the clamp loader variant in the host cell. *Inset*: Method to generate T4^del^ from the wildtype T4 bacteriophage using CRISPR-based genome engineering. (C) Schematic diagram showing the construction of chimeric clamp-loader variants. (D) Relative fitness of the chimeric clamp-loader variants from the phage-propagation assay.

Our work uses T4 bacteriophage as a model system for understanding the mutational response of the DNA replication machinery. The T4 phage genome encodes a complete set of replication proteins, including a replicative DNA polymerase and a clamp loader/clamp system. The pentameric T4 clamp-loader complex comprises of four identical ATPase subunits (gp44, denoted B-E in Fig. 1A), encoded by gene 44, and the clasp subunit (gp62, subunit A in Fig. 1A) encoded by gene 62 (Kelch et al. 2011). Clamp-loader subunits consist of N-terminal AAA+ modules and C-terminal collar domains that are responsible for maintaining the pentameric assembly (Guenther et al. 1997; Jeruzalmi et al. 2001; Bowman et al. 2004; Kelch et al. 2011). ATP binds at interfacial sites between neighboring AAA+ modules. The binding of primer-template DNA in a central channel formed by the AAA+ modules and the sliding clamp triggers the hydrolysis of ATP and the dissociation of the clamp loader from the sliding clamp, resulting in release of the clamp around DNA (Simonetta et al. 2009; Marzahn et al. 2014; Liu et al. 2017; Kelch et al. 2012; Zheng et al. 2022).

We developed a high-throughput T4 phage-propagation assay to measure the fitness of replication genes in their natural context (Fig. 1B) (Subramanian et al. 2021). The phage-propagation assay utilizes a genetically modified T4 phage, referred to as T4^del^, from which the genes encoding the sliding clamp (gene 45) and the clamp-loader subunits (genes 44 and 62) have been deleted (Fig. 1B, inset). Although T4^del^ phage particles can infect *E. coli* cells successfully, the phage genome cannot be replicated because of the missing components in the replication machinery. In the assay, T4^del^ is used to infect an *E. coli* cell library in which each cell carries plasmid-borne variants of the missing replication genes. Upon phage infection, variant genes from the plasmid are copied into the genome of T4^del^ using a CRISPR-Cas12a system. The number of phage particles produced from a bacterial cell is proportional to the function of the variant genes expressed in that cell. By measuring the frequency of each variant gene in the population using Illumina sequencing – in the plasmid library before phage infection and in the recombined phage population after infection – the fitness of each variant can be calculated.

We used the phage-propagation assay to perform a deep mutational scan of the clamp-loader complex and found that the clamp loader exhibits high mutational tolerance across most of its structure (Subramanian et al. 2021). Except for a sparse set of spatially contiguous residues in the functional core of the protein that are involved in ATP binding and hydrolysis and DNA recognition (constituting ∼10% of the clamp-loader complex), most residues in the clamp-loader complex tolerate point mutations without substantial loss in fitness. The mutational data also showed that the fitness of the wildtype clamp loader is not improved further by point mutations, suggesting that under normal conditions of bacterial growth, phage propagation is not limited by clamp-loader function (Subramanian et al. 2021).

An important question in understanding adaptive processes in molecular evolution concerns the capacity of proteins to adapt to changes in their environment, or to recover from mutational damage. We investigate the adaptive capacity of a defective variant of the clamp loader in terms of its ability to rescue the functional defect during phage propagation. The variant clamp loader, which is a chimera created by replacing the AAA+ modules of the T4 clamp loader by the corresponding modules from another clamp loader, is barely able to support phage replication and thus provides a sensitized genetic background that enable us to identify gain-of-function mutations. We subjected the phage bearing the chimeric complex to directed evolution and saturation point mutagenesis and identified mutations that compensate for the decreased fitness of the chimeric clamp-loader variant.

## Results and Discussion

### Engineering a clamp-loader variant with substantially reduced fitness

The structures of the AAA+ modules of clamp loaders are conserved across all branches of life, but their sequences have diverged considerably. The sequence of the AAA+ modules in bacteriophage clamp loaders have pairwise sequence identities as low as 25% (based on the alignment reported in (Subramanian et al. 2021)). In eukaryotic clamp loaders (RFC complex), each subunit of the pentameric assembly is encoded by a different gene. This has allowed the AAA+ modules in each subunit to further specialize in function and diverge in sequence (Schmidt et al. 2001). The pairwise sequence identities between the AAA+ module of the T4 clamp loader and each of the AAA+ modules of the yeast RFC complex range from 19% to 26% (Supp. Fig. 1A).

To generate a variety of clamp loaders with different degrees of loss of fitness, we created chimeric clamp-loaders in which the AAA+ module of the ATPase subunit of the T4 clamp loader (residues 1 to 230) was replaced by corresponding modules from other clamp loaders (Fig. 1C). We then used the phage-propagation assay to rank-order the chimeric variants by fitness. The chimeric clamp-loader variants generated in this fashion utilize the sliding clamp and other components of the replication system from T4 phage. Therefore, any reduction in fitness that result from propagating phage using the chimeric variants are due to non-optimal interactions between the transplanted AAA+ modules and the rest of the T4 replication machinery, or due to inefficiencies intrinsic to the AAA+ module itself.

Our creation of these chimeric clamp loaders is an artificial strategy that decreases fitness by swapping in catalytic domains from a different clamp loader. We note that natural recombination events within phage genomes can potentially yield clamp loaders with chimeric configurations, akin to the variants discussed above. When two different phage particles infect a host cell, the genomes of the progeny are mosaics of the parents due to extensive recombination (Hatfull 2008; Hendrix et al. 1999). While this is possible, the work presented here is motivated by an interest in studying how protein systems adapt in response to a reduction in fitness, rather than a specific interest in these chimeric systems.

We tested five chimeric clamp-loader variants using the phage-propagation assay. Two variants were generated by using AAA+ modules from bacteriophage clamp loaders: one from RB69 phage (83% sequence identity to the T4 AAA+ module) and one from *Aeromonas salmonicida* phage 44RR2.8t (denoted RR2 phage (Nolan et al. 2006; Petrov et al. 2006), with 64% sequence identity to the T4 AAA+ module) (Supp. Fig. 1A). Three chimeric variants were generated with AAA+ modules from a eukaryotic system (the yeast Replication Factor C, or RFC complex), each having <30% sequence identity to the T4 AAA+ module (Supp. Fig. 1A).

We used the phage-propagation assay to measure the ability of the chimeric clamp-loader variants to support replication of T4 phage (Fig. 1B). We calculate the relative fitness (*F*) of each variant in the assay as follows:

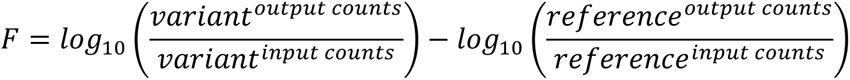

where *F* is the relative fitness score for the variant relative to the reference (T4 clamp loader in this experiment). The *output counts* correspond to the number of counts observed in the recombinant-phage library for the variant and reference clamp loader sequences. The *input counts* correspond to the number of counts observed in the starting plasmid library for the variant and reference. A variant with a relative fitness score of zero propagates at the same rate as the wildtype clamp loader. Variants with fitness scores of –1 and –2 propagate 10-fold and 100-fold slower, respectively, than the reference.

We used the phage-propagation assay to measure the relative fitness of the chimeric clamp-loader variants (Fig. 1D). The three variants with yeast RFC AAA+ modules are essentially non-functional, with relative fitness scores that are comparable to that for the null variant (a variant in which the three codons following the start codon of gene 44 are replaced with stop codons). Chimeric variants generated using AAA+ modules from RB69 and RR2 phage clamp loaders have fitness scores that are several orders of magnitude lower than that of the wildtype clamp loader, but are still measurable (the propagation rates are ∼5x10^4^ and ∼5x10^3^ fold lower than wildtype, respectively). In a separate experiment where the wildtype T4 phage was competed with T4^del^ recombined with the RR2-chimeric clamp loader, a similar fitness difference between the two variants was observed (data not shown). This suggests that the influence of variant clamp loader on the CRISPR-mediated recombination can be disregarded.

To identify mutations in chimeric clamp loaders that result in increased fitness, we focused on the variant with the AAA+ module from RR2 phage (the RR2/T4 chimera) since it is more divergent in sequence compared to the AAA+ module from RB69 phage (64% vs 83% sequence identity). Of the 230 residues in the AAA+ module, 83 are different between the RR2 and T4 sequences (Supp. Fig. 2A). Using previously determined mutational data for the T4 clamp loader (Subramanian et al. 2021), we compared the distribution of fitness effects for each of these 83 substitutions to the T4 clamp loader sequence, taken individually, with the distribution of all possible single amino acid substitutions in the T4 background (Supp. Fig. 2B). The analysis shows that 80 of the 83 substitutions, when made individually in the wildtype context, have a marginal effect (less than three-fold) on T4 phage propagation. Three substitutions (H44N, G209N, S213F) result in a moderate (∼10-fold) reduction in phage propagation. Mutations of Gly 86, which we focus on in the discussion below, have only mild effects (less than three-fold decrease in phage propagation) when introduced in the context of the wildtype clamp loader (Supp. Fig. 3).

### Directed evolution to recover fitness in the RR2/T4 chimeric clamp loader

We modified the phage-propagation platform to carry out directed-evolution experiments on the RR2/T4 chimeric clamp loader. We used the CRISPR-Cas12a genome editing system (Zetsche et al. 2015) to generate a new strain of T4 phage – referred to as T4^pol_del^ phage – in which the genes encoding the T4 DNA polymerase (gene 43) and the T4 clamp loader (genes 44 and 62) are deleted (Fig. 2A; for details, see Methods). We then inserted genes encoding the RR2/T4 chimera into T4^pol_del^ phage to generate T4^chimera^ phage (Fig. 2A). The T4^chimera^ phage lack a gene for DNA polymerase and can only propagate in host cells harboring a plasmid-borne copy of the T4 DNA polymerase gene.

**Figure 2.**
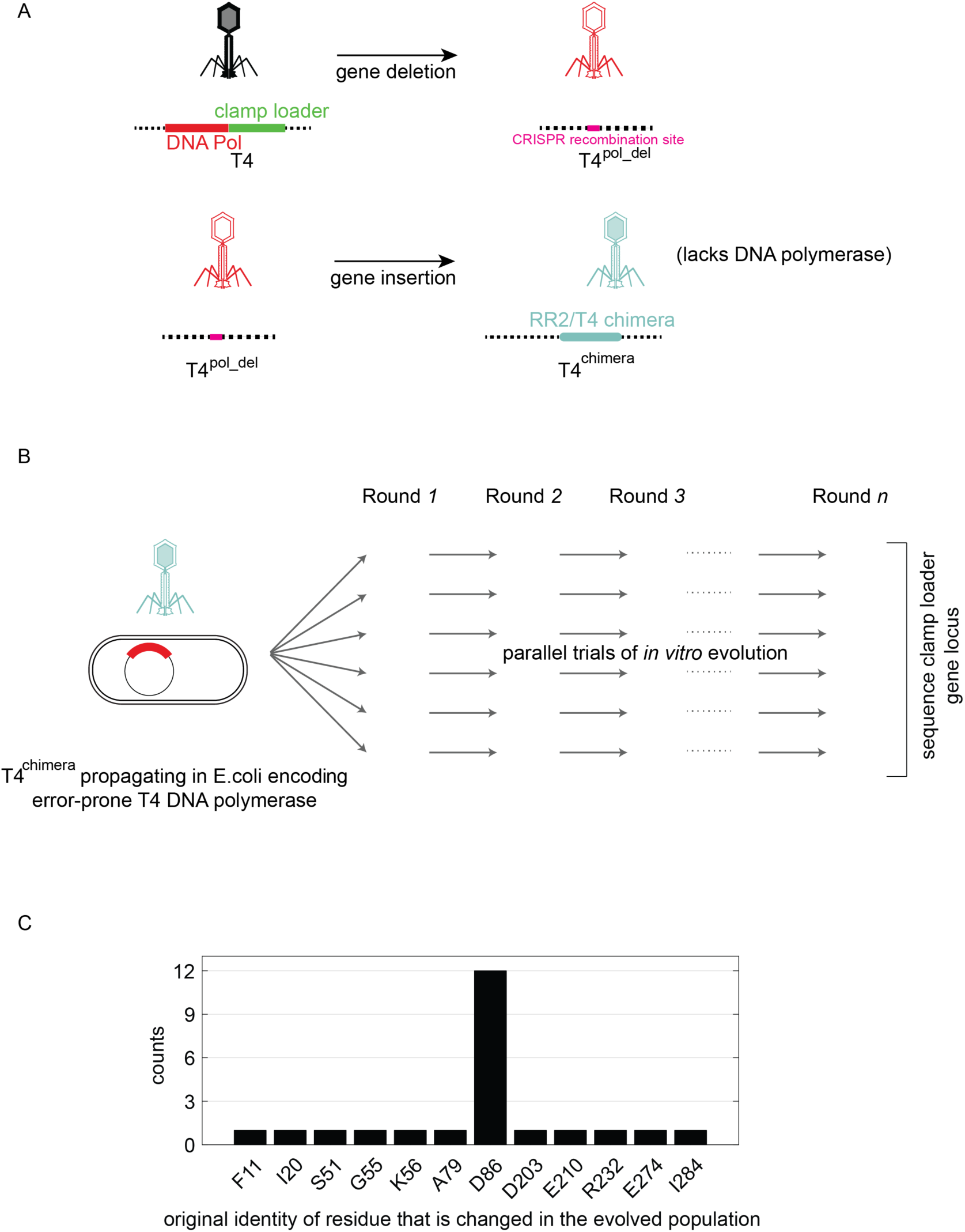
Platform for performing *in vitro* evolution to improve the fitness of bacteriophage T4 propagating using the RR2/T4 chimeric clamp loader. (A) Schematic diagram depicting the construction of the T4^chimera^ used to perform *in vitro* evolution. (B) Schematic representation of the *in vitro* evolution protocol used in the study. (C) Mutations identified by subjecting the T4^chimera^, bearing the RR2/T4 chimeric clamp loader, to *in vitro* evolution.

When propagated, T4^chimera^ phage produce new T4^chimera^ phage particles (devoid of the DNA polymerase gene) in numbers that are proportional to the activity of the chimeric clamp-loader variant. In order to perform *in vitro* evolution, T4^chimera^ phage are propagated over many rounds (10-15 rounds in this study) so that individual phage genomes that spontaneously acquire beneficial mutations get enriched in the phage population. The rate of finding beneficial mutations can be accelerated by increasing the overall mutation rate, through the use of error-prone variants of T4 DNA polymerase (Langhorst et al. 2012). We used a single-mutant variant of the T4 DNA polymerase (D219A), with a ∼1,000-fold higher mutation rate than the wildtype polymerase (Frey et al, 1993), to passage phage through multiple rounds of *in vitro* evolution.

We carried out 15 independent trials of *in vitro* evolution with the error-prone variant of T4 DNA polymerase. For each trial, after 10 rounds of passaging phage, we sequenced the genes encoding the clamp loader and the sliding clamp in the phage population using Sanger sequencing and identified the dominant mutations. A total of 23 point mutations were identified in this way. Strikingly, 12 of the 15 trials identified mutations to a single residue, Asp 86, in the AAA+ module of the RR2/T4 chimera, with all other mutations observed in only one of the trials (Fig. 2B and Supp. Fig 4A).

We observed that substitutions at residue Asp 86 of the AAA+ module of the chimeric clamp loader first appear in the population at different rounds, but once they appear, they immediately take over the population in the subsequent round of phage propagation (representative sequencing data are shown in Supp. Fig. 4B, 4C), suggesting that these mutations result in a substantial increase in the fitness of the T4^chimera^.

### The fitness effects of Asp 86 substitutions

The Sanger sequencing covers only the genetic locus of the clamp loader and the sliding clamp, and so will miss gain-of-function mutations that occur elsewhere on the genome. Therefore, we verified that the substitutions at Asp 86 in the RR2/T4 chimeric clamp-loader do indeed result in gain of fitness on their own. We carried out saturation mutagenesis of position 86 of the RR2/T4 chimera and tested all variants using the phage-propagation assay (Fig. 3A). Substitutions of Asp 86 by glycine, alanine and asparagine residues – which appeared in the *in vitro* evolution experiment – result in 10-300-fold increases in phage propagation with respect to the RR2/T4 chimera (Fig. 3A). Importantly, the saturation mutagenesis data show that several other substitutions of Asp 86 that remove the negatively charged sidechain of Asp 86, such as substitutions by arginine, histidine, serine, cysteine, methionine, tyrosine and phenylalanine, also increase the fitness of the RR2/T4 chimera.

**Figure 3.**
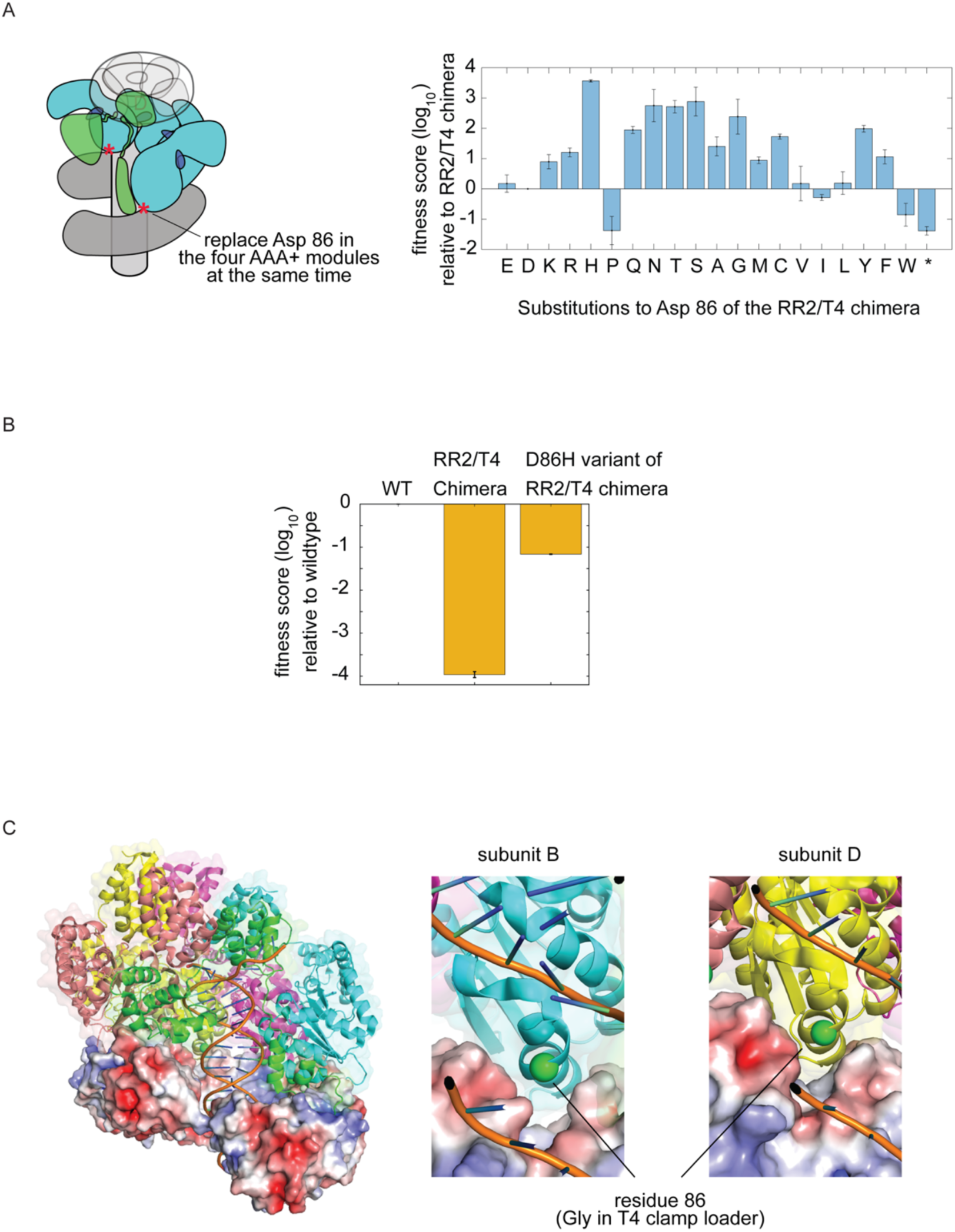
Fitness effect of substituting Asp 86 of the AAA+ module in the RR2/T4 chimeric clamp loader. (A) Deep mutational scan of position 86 of the AAA+ module in the RR2/T4 chimeric clamp loader. (B) Fitness scores of the RR2/T4 chimeric clamp loader and its D86H variant, relative to the wildtype T4 clamp loader. (C) Location of residue 86 of the AAA+ module in the structure of the clamp loader bound to the sliding clamp and DNA (PDB ID: 3U60). The surface rendering of the sliding clamp is colored by electostatics potential, calculated with the Adaptive Poisson-Boltzmann Solver (APBS), ranging from –5.0 kT (red) to +5.0 kT (blue).

We note that the D86H mutation, identified by saturation mutagenesis as causing the largest increase in fitness for the chimera, was not observed in the 15 independent trials of directed evolution. We suspect that the failure of the directed evolution experiment to identify the importance of the D86H mutation is a result of mutational bias in the error-prone polymerase employed.

The histidine substitution at position 86 results in the most substantial increase in the fitness of phage (by ∼3,000-fold) (Fig. 3A). In order to compare the fitness of the D86H variant of the RR2/T4 chimera with that of the wildtype T4 clamp loader, we performed a phage-propagation assay competing three clamp loader variants: the wildtype T4 clamp loader, the RR2/T4 chimera, and the D86H variant of the RR2/T4 chimera (Fig. 3B). The conditions of the experiment are such that there is only one wave of phage infection, with very little cells uninfected cells at the onset of the experiment (multiplicity of infection is 3). In these conditions, the RR2/T4 chimera propagates at a ∼10,000-fold slower than the T4 clamp loader. The fitness score of the D86H variant of the RR2/T4 chimera is –1.2 relative to the wildtype T4 clamp loader, which corresponds to ∼20-fold slower propagation compared to the wildtype variant. Thus, a single point mutation (D86H) has repaired much of the deficiency of the RR2/T4 chimeric clamp-loader.

The ability of several different substitutions of Asp 86 to rescue fitness suggests that the aspartate residue at this position in the RR2/T4 clamp loader may be involved in unfavorable interactions in the context of the chimeric clamp-loader complex, and that the increased fitness is a consequence of removing these unfavorable interactions. Residue 86 is part of helix α4, which along with helices α5 and α6 constitute the “central coupler” in AAA+ assemblies (Subramanian et al. 2021). The central coupler is a structurally contiguous unit that spans the AAA+ modules within the assembly, coupling the ATP-binding sites to DNA, to adjacent subunits, and to the sliding clamp. Helix α4 contacts the surface of the sliding clamp.

Inspecting the structure of the DNA-bound form of the T4 clamp loader (Kelch et al. 2011) (Fig. 3C) provides a possible explanation for why the replacement of Asp 86 in RR2/T4 chimera results in an increased fitness. For two of the four AAA+ modules in the structure, at positions B and D of the clamp-loader complex (Fig. 3C), residue 86 – glycine in the T4 clamp loader – is positioned just above a region of negative electrostatic potential on the surface of the T4 sliding clamp. This suggests that replacing the T4 AAA+ module with that of RR2 phage, which has the negatively-charged sidechain of Asp 86 located at the interface with the clamp, would introduce an unfavorable electrostatic interaction, and a decrease in the fitness of the clamp loader.

That Asp 86 causes unfavorable electrostatic interactions between the RR2/T4 chimeric clamp loader and the T4 sliding clamp is consistent with mutagenesis data for the T4 sliding clamp, as discussed below. The structures also suggest an additional explanation. Asp 86 is located next to Arg 85, a residue that is extremely sensitive to mutation in the phage-propagation assay and highly conserved (Subramanian et al. 2021). The structure of the DNA-bound clamp-loader complex shows that Arg 85 contacts the phosphate backbone of DNA (Kelch et al. 2011). We modeled the structure of the AAA+ module of the RR2 clamp loader using Alphafold (Mirdita et al. 2022; Jumper et al. 2021) and used it to carry out molecular dynamics simulations (six independent trajectories, 2 μs each) of the free AAA+ module in the absence of DNA and the sliding clamp. When not able to interact with DNA, the simulations suggest that Arg 85 can make a stable ion pair with Asp 86 (Supp. Fig. 5A, 5B). It is possible that this interaction is also present in the context of the RR2/T4 chimera and the T4 sliding clamp, thereby reducing DNA affinity by competing with DNA for Arg 85, but this possibility awaits verification by additional experiments.

### Deep mutational scan of the sliding clamp

The idea that electrostatic repulsion between the clamp loader and the sliding clamp underlies the reduction in fitness of the RR2/T4 chimeric clamp-loader is supported by the results of saturation mutagenesis of the T4 sliding clamp (Supp. Fig. 6). In this experiment, the T4 sliding clamp is subjected to saturation single-point mutagenesis, and phage propagation using the RR2/T4 chimeric clamp loader is measured. The fitness scores for the sliding clamp mutants are calculated relative to phage propagating with the unmutated T4 sliding clamp and the RR2/T4 chimeric clamp-loader. Residues Glu 156 and Asp 157 of the T4 sliding clamp contribute to the negative electrostatic potential on the surface of the clamp in the region adjacent to where the helix bearing Asp 86 interacts with the clamp (Fig. 4A). We observe that mutating these two residues increases fitness (Fig. 4B), with the E156T and D157S variants resulting in fitness scores of 1.6 ±0.6 (∼10- fold faster phage propagation than that for the RR2/T4 chimera) and 2.5 ±0.1 (∼300-fold faster phage propagation than that for the RR2/T4 chimera), respectively.

**Figure 4.**
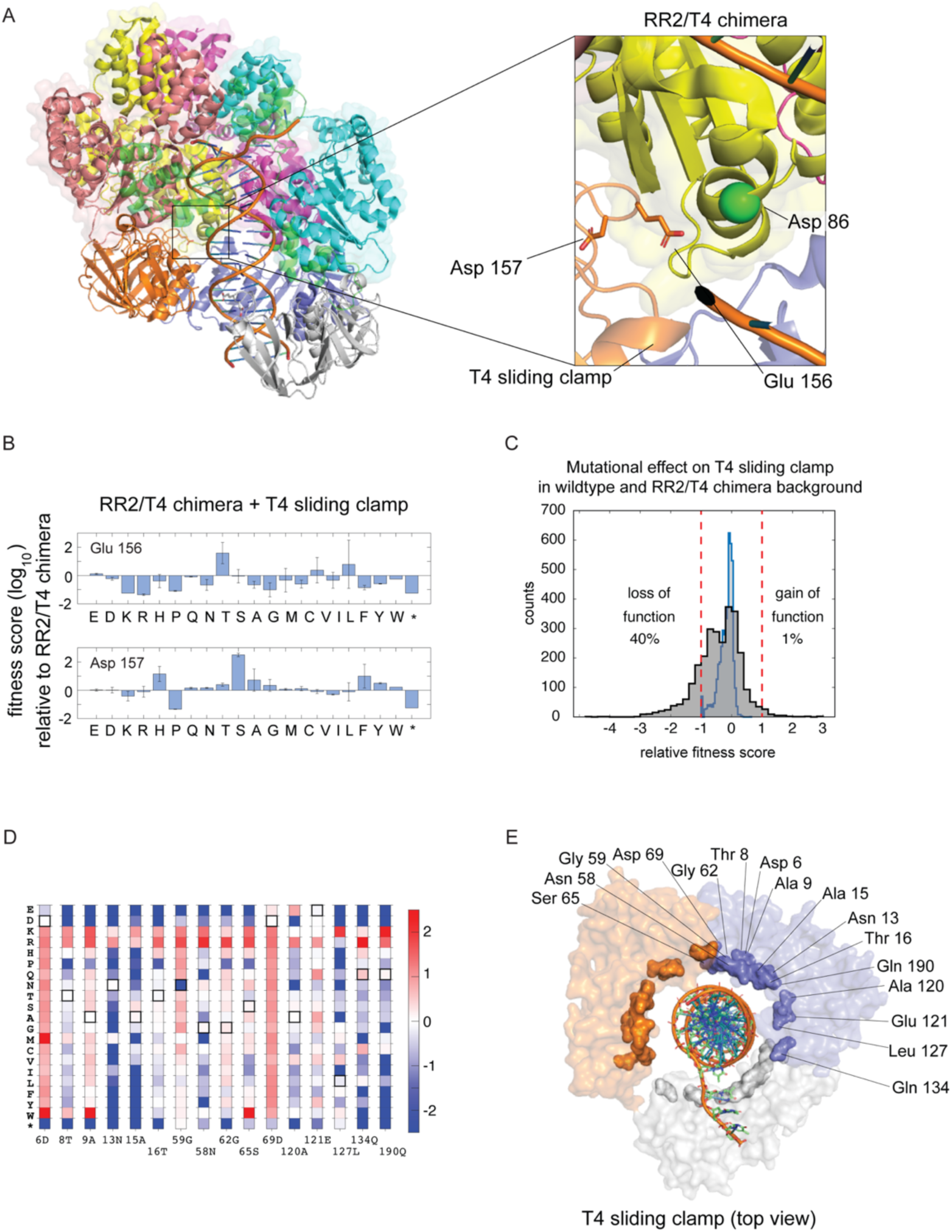
Deep mutational scan of the T4 sliding clamp in the context of the RR2/T4 chimeric clamp loader. (A) Acidic residues of the T4 sliding clamp that could interact unfavorably with Asp 86 of the AAA+ module in the RR2/T4 chimeric clamp loader. (B) Fitness scores for the mutational scan of the acidic residues in A. (C) Distribution of mutational effects of the T4 sliding clamp in the context of the wildtype T4 clamp loader (blue) and the RR2/T4 chimeric clamp loader (black). (D) Mutational profiles of residues in the T4 sliding clamp containing mutations that improve the fitness, by at least 10-fold, of T4 phage propagating with the RR2/T4 chimeric clamp loader. (E) Residues in D represented on the structure of the T4 sliding clamp (PDB ID: 3U60).

The data show that ∼1% (43 of 4313) of the mutations to the T4 sliding clamp increase the rate of phage propagation by more than 10-fold relative to that with the RR2/T4 clamp loader (Fig. 4C). These gain-of-function mutations occur at 16 positions in the clamp, the mutational profiles of which are represented as a heatmap in Fig. 4D. The heatmap reveals a striking pattern: substitutions at these positions to arginine or lysine, residues that have positively-charged sidechains, increases the fitness of phage. In fact, any substitution that removes the negatively-charged sidechain of Asp 6 and Asp 69 cause a fitness increase.

An explanation for this pattern emerges upon inspecting where these residues are located in the structure of the clamp loader in complex with DNA and the sliding camp (Kelch et al. 2011) (Fig. 4E). The 16 residues for which arginine and/or lysine substitutions increase fitness are proximal to DNA, and are located on the inner surface of the T4 sliding clamp. Introducing positively-charged residues at these positions can result in a favorable interaction with DNA. A multiple sequence alignment of ∼1,000 diverse sequences of the sliding clamp (Supp. Fig. 7) showed that arginine and lysine residues are sparsely sampled at these 16 positions. This suggests that the ability of the T4 clamp to increase the fitness of phage through DNA interactions mediated by these regions represents an evolutionarily underutilized route to increase fitness, at least among extant sliding clamps.

### Deep mutational scan of the clasp subunit in the RR2/T4 chimeric clamp loader

From a structural perspective, the clasp (A) subunit plays a crucial role in clamp loader function because it bridges ATPase subunits B and E, and plays a key role in recognizing the primer-template junction (Fig. 1A) (Kelch et al. 2011). The clasp straddles the two domains of the sliding clamp that move apart when the clamp opens. Given its central role in the structure, it is surprising that the clasp subunit is highly tolerant to mutations in the context of the wildtype T4 system (Subramanian et al. 2021). This highlights the fact that when the clamp loader is operating under optimal conditions, it is most sensitive to mutations that break the integrity of the essential catalytic machinery.

In the RR2/T4 chimeric clamp-loader, the clasp subunit is that of the T4 system. We reasoned that the reduced fitness of the RR2/T4 chimera would sensitize the clasp to mutations. We performed a deep mutational scan of the clasp subunit in the context of the RR2/T4 chimera and indeed observed a pattern of mutational sensitivity that is consistent with expected structural importance of the clasp subunit (Supp. Fig. 8).

We highlight the mutational sensitivity of the clasp using a representative region of the clasp subunit, the N-terminal arm (residues 2-44, Fig. 5A). This segment of the clasp subunit contacts the sliding clamp near the location of the opening in the sliding clamp. It also interacts with the B-subunit of the clamp-loader complex, and is situated adjacent to DNA. The importance of the structural integrity of the clasp subunit is highlighted by the fact that proline substitutions in helical regions, but not in loops, reduce the fitness of the RR2/T4 chimera (Fig. 5B, Supp. Fig. 8). Proteins that interact with the sliding clamp, including clamp loaders and DNA polymerases, engage the clamp using a conserved binding motif, the PCNA interacting protein-box or the PIP-box (Warbrick 1998; Gulbis et al. 1996). In T4 phage, the principal interaction with the clamp utilizes a PIP-box in the clasp subunit (Leu 3 and Phe 4) (Fig. 5A). The data show that mutations to Leu 3 and Phe 4 substantially reduce (by more than 10-fold) the fitness of the RR2/T4 chimera (Fig. 5B).

**Figure 5.**
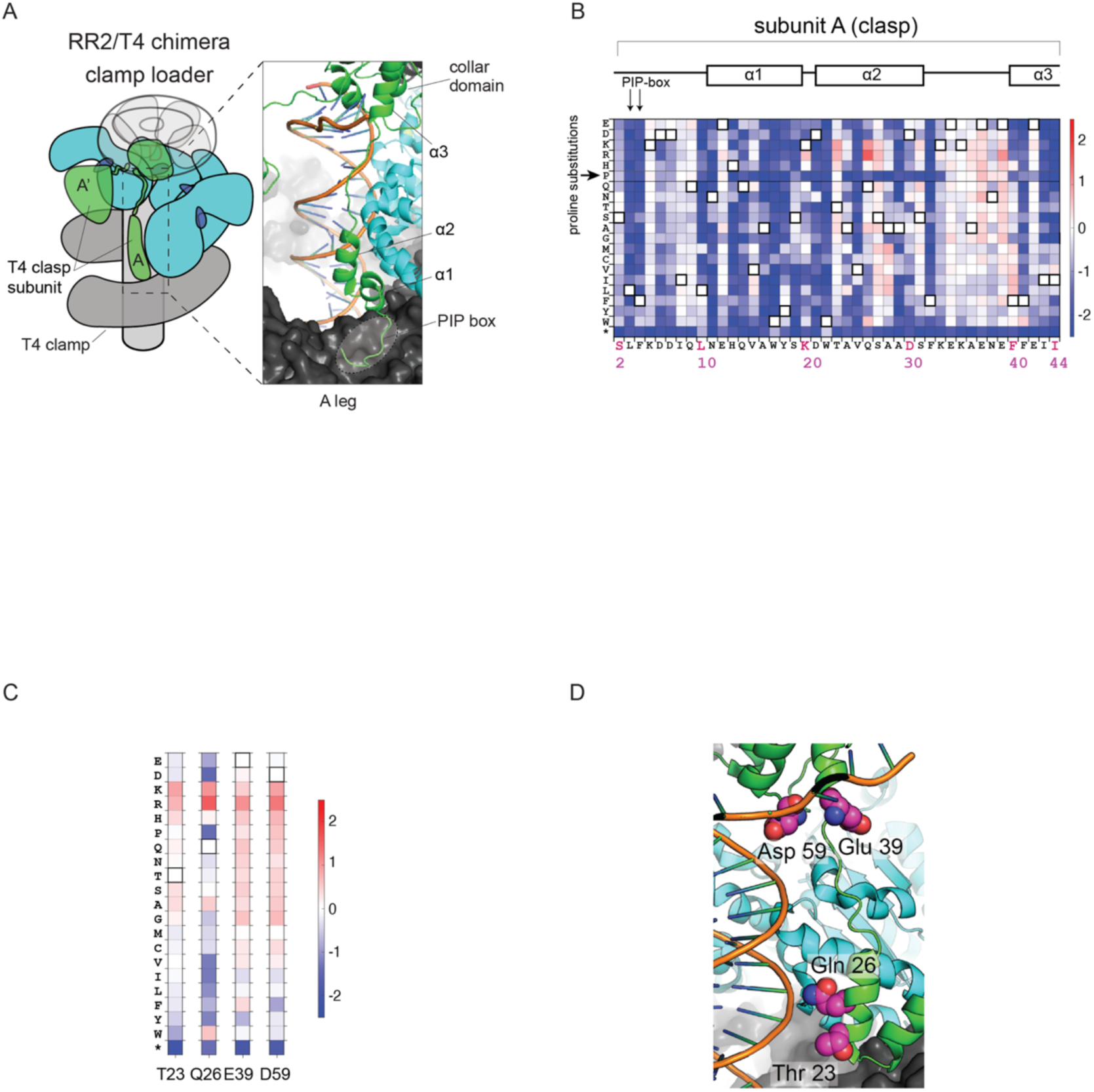
Deep mutational scan of the T4 clasp subunit in the context of the RR2/T4 chimeric clamp loader. (A) The N-terminal arm of the T4 clasp subunit contacts the sliding clamp, DNA and the B-subunit ATPase of the clamp loader. (B) Deep mutational scan of the N-terminal arm of the T4 clasp subunit. (C) Mutational profiles of residues in the T4 clasp subunit containing mutations that improve the fitness, by at least 10-fold, of T4 phage propagating with the RR2/T4 chimeric clamp loader. (D) Residues in C represented on the structure (PDB ID: 3U60).

Saturation mutagenesis of the clasp subunit identifies eight gain-of-function substitutions that result in a greater than 10-fold increase in phage propagation in the context of the RR2/T4 chimeric clamp loader. These substitutions occur in five residues in the clasp (Thr 23, Gln 26, Glu 39, Asp 59 and Asn 140) (Fig. 5C). Except for Asn 140, the other four residues are proximal to DNA in the ternary structure of the clamp loader bound to DNA and the sliding clamp (Fig. 5D). The mutational profile at these four positions in the RR2/T4 chimera shows that introducing arginine or lysine, which can increase DNA affinity by interactions with the phosphate backbone, increases the rate of phage propagation (Fig. 5C). Thus, both the clasp subunit and the sliding clamp possess the ability to acquire single mutations that can increase DNA affinity. The fifth residue that shows gain-of-function in the clasp – Asn 140 – is a non-interfacial residue, and the mechanism underlying the 20-fold increase in phage propagation caused by the N140T mutation is unclear at present.

### Deep mutational scan of the AAA+ module of the RR2/T4 chimeric clamp loader

We performed a deep mutational scan of the AAA+ module in the RR2/T4 chimeric clamp loader to identify mutations to the RR2 AAA+ module that are beneficial in the chimeric context (Supp. Fig. 9). The data show that nine positions (Pro 50, Asp 82, Asp 86, Cys 98, Val 106, Ala 112, Gly 113, Ala 115 and Gly 223), have at least two mutations that can cause a >10-fold increase in fitness. These nine residues can be grouped into three sets (Supp. Fig. 10). The first set of residues (Asp 86, discussed earlier, and Cys 98) are on helix α4 and interact with the sliding clamp, and substitutions to this set of residues are likely to improve fitness by increasing the affinity of the chimeric clamp-loader for the clamp. The second set of residues (Pro 50 and Gly 223) belong to a group of residues reported in another study of the T4 clamp loader (*Marcus, Huang et al, in press*) that destabilize the DNA-free inactive structure, facilitating the transition to the DNA-bound active state. The third set of residues (Asp 82, Val 106, Ala 112, Gly 113, Ala 115) contact DNA-sensing residues (Lys 80 and Arg 111). Substitutions to this set of residues likely causes a re-positioning of the DNA-sensing residues that increases DNA affinity and thereby, the fitness of phage.

### Conditional neutrality in the RR2/T4 chimeric clamp loader

Conditional neutrality is a major emerging theme in protein adaptation that refers to mutations that are neutral in one setting but confer a fitness benefit in a new setting (Wagner 2008; Hayden et al. 2011; Zheng et al. 2019; Draghi and Plotkin 2011). The deep mutational scan of the T4 clamp loader complex revealed many fitness-neutral mutations (Subramanian et al. 2021). We analyzed the results from deep mutational scanning of the RR2/T4 chimeric clamp loader to identify conditionally neutral mutations: mutations that are beneficial in the chimeric context and neutral in the context of the wildtype T4 clamp loader (Fig. 6). Mutations were deemed neutral in the wildtype T4 clamp-loader setting if their effect on fitness was within three standard deviations from the mean of the fitness distribution of the synonymous variants of the wildtype T4 sequence. Using this threshold, the neutral range of fitness was within 4-fold of the wildtype for the sliding clamp (Supp. Fig. 11A), within 3-fold of the wildtype for the clasp (Supp. Fig. 11B), and within 5-fold of the wildtype for the AAA+ module (Supp. Fig. 11C). Mutations in the RR2/T4 chimeric clamp-loader setting were considered beneficial if they cause at least a 10-fold increase in fitness relative to the fitness of the RR2/T4 chimera. We considered all positions in the AAA+ module in this analysis, irrespective of whether the positions are identical or different between the wildtype T4 and chimeric clamp loaders. Using these thresholds, all the beneficial mutations – 53 mutations in the T4 sliding clamp (Fig. 6A), 6 mutations in the T4 clasp (Fig. 6B) and 33 mutations to the AAA+ module (Fig. 6C), are conditionally neutral.

**Figure 6.**
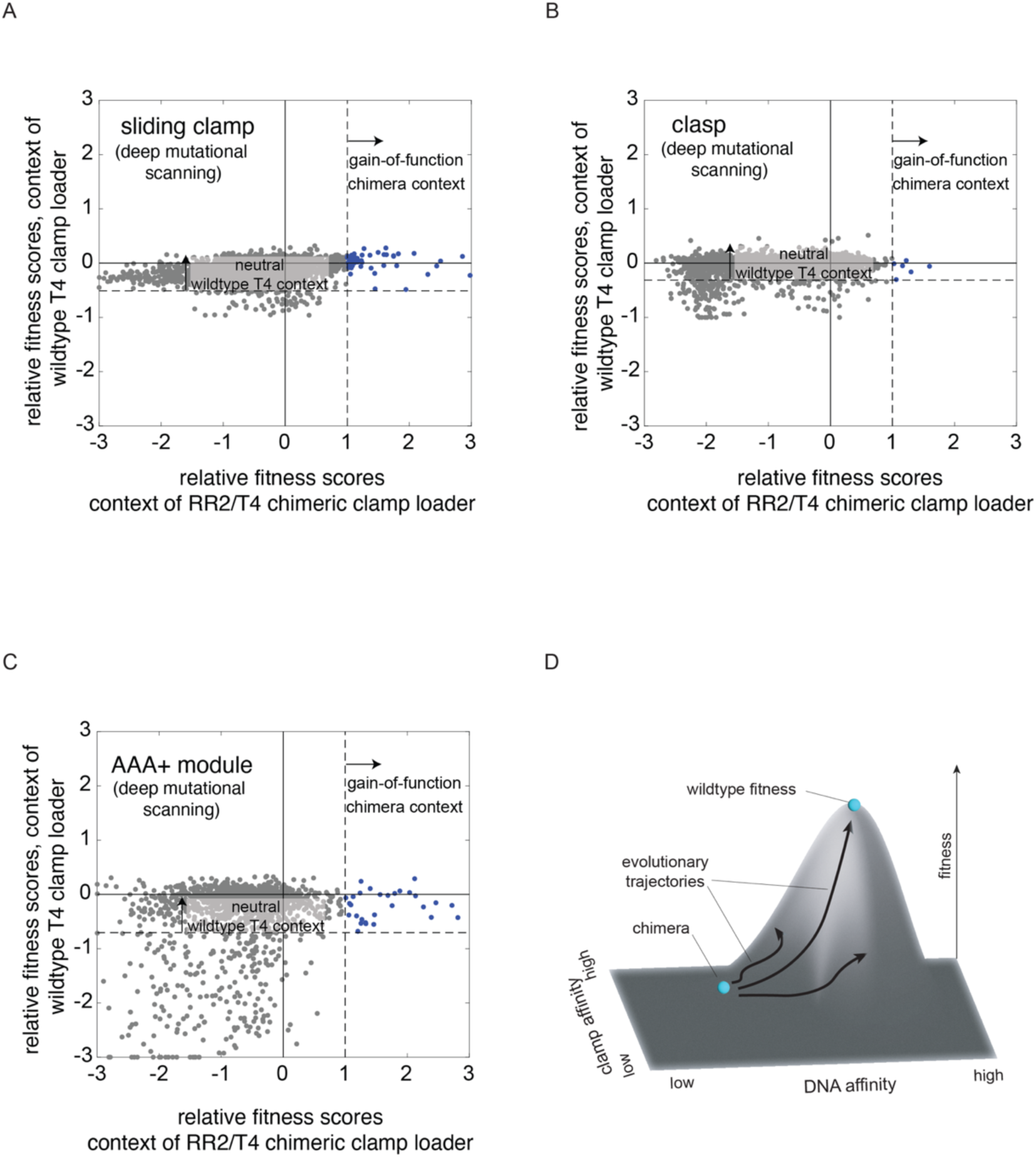
Conditionally neutral mutations in the clamp-loader complex. Scatter plots comparing the fitness effect of mutations in the context of the RR2/T4 chimeric clamp loader and the wildtype T4 clamp loader, for the sliding clamp (A), the clasp subunit (B) and the AAA+ module (C). (D) Schema depicting the idea that clamp-loader function can be optimized along many properties (DNA binding and clamp binding shown), but evolution balances these properties for optimal fitness for a given condition, say for T4 bacteriophage propagating in *E. coli*.

### Concluding Remarks

In this work, we studied the adaptive capacity of bacteriophage DNA polymerase clamp loaders by using a defective variant of the T4 clamp loader in which the AAA+ modules are replaced by those from a clamp loader in RR2 phage. The resultant chimeric clamp loader causes the T4 phage propagation rate to drop by ∼5000-fold. We subjected the T4-RR2 chimeric phage to directed evolution and found that point mutations to a single residue – Asp 86 – in the RR2 AAA+ module can dramatically increase phage fitness. Based on the structure of the T4 clamp-loader complex, it appears that mutation of Asp 86 repairs the chimeric clamp loader by reducing electrostatic repulsion with the T4 sliding clamp. A deep mutational scan of the T4 sliding clamp, the T4 clasp subunit and the swapped-in RR2 AAA+ module, carried out in the context of the chimeric clamp loader, revealed that substitutions to residues proximal to DNA increase the fitness of phage replicating with the chimeric clamp loader, presumably by tuning DNA interaction. Taken together, these results show how natural selection can exploit the adaptive potential latent in the clamp loader subunits to improve fitness through single amino acid changes.

In a parallel study (*Marcus, Huang et al, in press*), we investigated the adaptive capacity of the clamp loader by starting from a T4 clamp loader with a single mutation (D110C) in the conserved DEAD box motif that resulted in a mild (∼6-fold) reduction in function. We mapped the mutational response of the AAA+ module of the mutant clamp loader and found that this mild defect is rescued by a diverse set of mutations that are distributed throughout the AAA+ module. The locations of the mutations map to regions of conformational change between the active DNA-bound state of the clamp loader and the newly identified DNA-free inactive states as shown by cryo-EM analysis (*Marcus, Huang et al, in press*). These mutations, which individually only have a mild effect on fitness, were only identified because the fitness deficiency due to D110C is itself very mild. In contrast, the much more substantial defect in the RR2/T4 chimeric clamp-loader is due to sub-optimal interactions between non-cognate components. Thus, the two studies highlight complementary mechanisms by which the clamp loader can alter its function – by modulating the strength of interaction between its components and its substrates (the sliding clamp and DNA) and by altering the free-energy balance between the active and inactive conformations of the complex.

Conditionally neutral mutations play a pivotal role in facilitating the evolutionary diversification of proteins since they can pre-exist as neutral, standing genetic variation, waiting to reveal their beneficial effects when the environmental context changes (Luria and Delbrück 1943; Draghi and Plotkin 2011; Dellus-Gur et al. 2013; Hayden et al. 2011; Payne and Wagner 2014; Raman et al. 2016; Jayaraman et al. 2022; Wagner 2005). For example, the conditionally neutral mutations in the T4 sliding clamp can persist in a population of T4 phage without affecting reproductive fitness, only to prove advantageous when chimeric clamp loaders arise due to genetic transfer between two phage genomes. We revealed conditional neutrality in the clamp loader by using an artificially generated chimeric variant that is defective by construction. An intriguing avenue for future work is to study the generality of the conditionally neutral mutations: are the mutations that are beneficial in the chimera also advantageous in a differential context? The fact that mutations at residue Pro 50 of the AAA+ module of the T4 clamp loader are beneficial in the chimeric clamp loader as well as in the genetic background of the D110C mutation (*Marcus, Huang et al, in press*) suggest that this might indeed be the true.

Biological systems have many properties that can be optimized to increase aspects of their function. For the clamp loaders, this could be DNA binding, ATP binding/hydrolysis, clamp binding or clamp release (Fig. 6D). Under a given set of conditions, evolution balances all these properties to maximize the fitness of the organism. Optimizing along just one property would reduce fitness – for example, a clamp loader that bound DNA too tightly would not release the clamp on DNA. In this study, we tipped the clamp loader off balance by creating chimeric clamp-loader variants that resulted in suboptimal inter-subunit interactions. Remarkably, natural selection can find a new balance between these properties rather quickly, with point mutations sufficing to get appreciable improvement in fitness.

### Key Resource Table

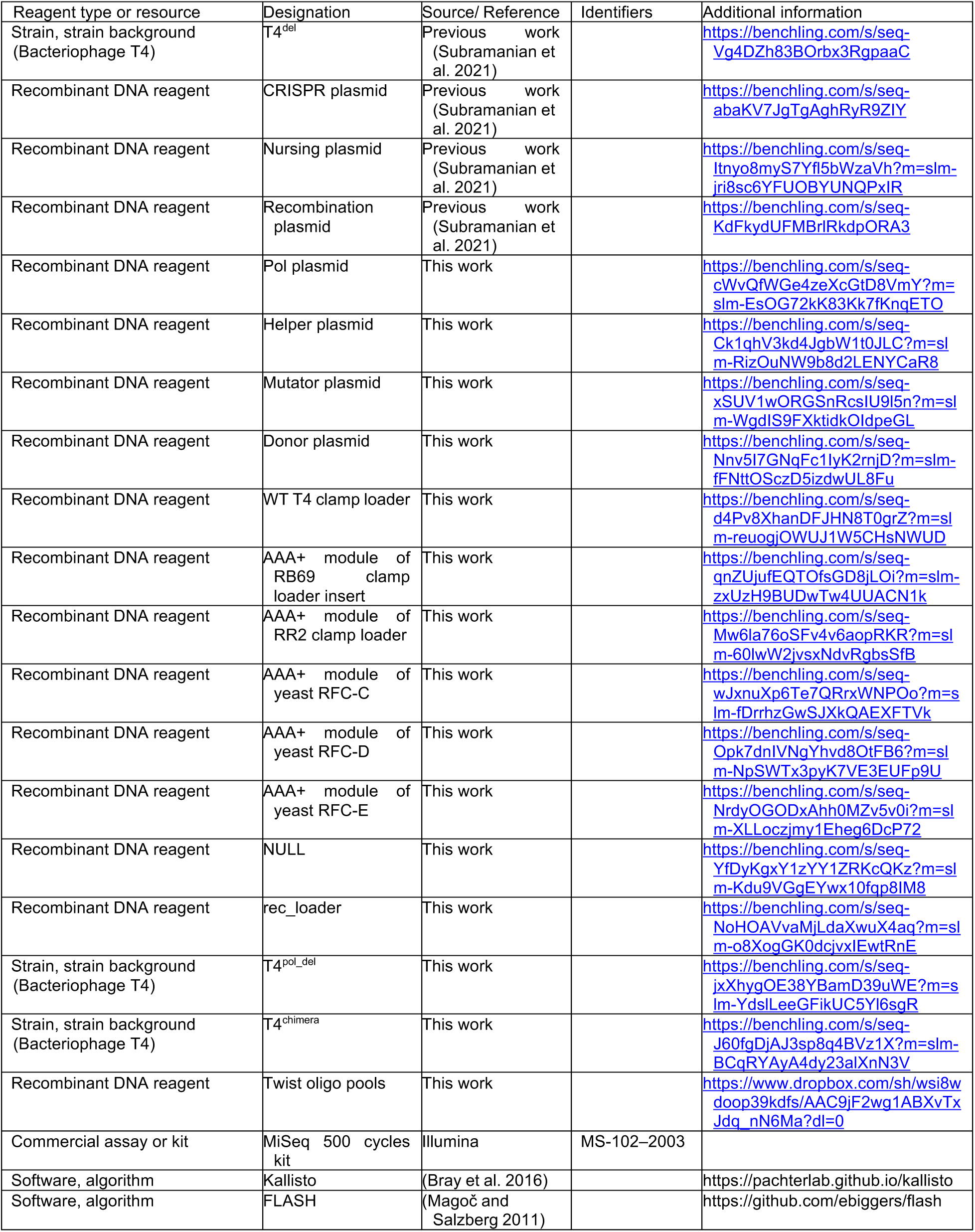

## Materials and methods

### Construction of chimeric variants of the clamp loader for the phage-propagation assay

The phage-propagation assay used to measure the ability of variant clamp loader variants to support phage replication depends on a recombination plasmid that carries the genes encoding the clamp-loader subunits (Subramanian et al. 2021). We had previously generated a recombination plasmid containing the genes for the T4 clamp loader complex, including gene 44 encoding the ATPase subunit and gene 62 encoding the clasp subunit (Subramanian et al. 2021). To generate chimeric variants of the clamp loader, we replaced residues 2-232 of the ATPase subunit of the T4 clamp loader with the corresponding residues from other AAA+ modules. DNA encoding other AAA+ modules were amplified from the genomes of *Saccharomyces cerevisiae*, bacteriophage RB69 and bacteriophage RR2. We introduced the BamHI restriction site (GGATCC) in gene 44, between regions encoding the AAA+ module and the collar domain for ease of cloning. We generated a NULL variant of the clamp loader by introducing three stop codons after the start codon of gene 44. Hyperlinks to the annotated plasmid sequences are provided in the key resource table.

### Construction of single-mutant libraries

We constructed comprehensive single-mutant libraries of the clamp, clasp, and AAA+ module of the chimeric clamp-loader complex using previously described methods (Subramanian et al. 2021). The primers used to generate the mutant library for the sliding clamp were re-used to generate the library in the recombination plasmid containing the chimera. Mutant libraries for the clasp subunit and the AAA+ module were generated using oligonucleotide pools ordered from Twist Biosciences (San Francisco, California). The AAA+ module library was generated using 10 pools, each encoding 480 sequences corresponding to 24 positions. The mutant library for the clasp subunit was generated using 3 pools, each encoding ∼1300 sequences corresponding to 65 positions. The sequences of the oligonucleotide pools are available here.

### Phage-propagation assay

The phage-propagation assay was performed as described in previous work (Subramanian et al. 2021). Briefly, the plasmid library encoding the RR2/T4 chimeric clamp-loader variants was transformed into bacterial cells containing the CRISPR plasmid and recovered in an overnight growth in selective media. ∼100 μl of the dense, overnight culture was inoculated into fresh ∼110 ml LB broth to start an exponentially growing culture. Plasmid was extracted from ∼10 ml of the culture (OD_600_ = 0.1) and the DNA was set aside as the starting library. Phage propagation was started by adding ∼10^8^ T4^del^ phage particles to the remainder of the culture (∼100 ml). This corresponds to a multiplicity of infection (MOI) of 0.1. The phage infection was allowed to proceed for ∼ 24 hrs at 37 °C in a shaking incubator, and then allowed to sit on the bench for the cell debris to settle. ∼1 ml of the culture was centrifuged to remove cell debris, and the supernatant was saved as the selected phage library. DNA samples were prepared for Illumina sequencing through three sequential PCR steps as described previously (Subramanian et al. 2021), and sequenced on the MiSeq sequencer using the 500 cycles kit.

### Analysis of the raw sequencing data

The different clamp-loader sequence variants present in the sequencing data were quantified using Kallisto version 0.48.0 obtained from GitHub. Kallisto index was built, using default parameters, from a custom fasta file containing all possible unique DNA sequences encoding the clamp loader variants. Relative fitness scores were calculated using custom MATLAB scripts available here.

### Generation of T4^chimera^ strain for directed evolution through phage propagation

The T4^chimera^ strain used in the directed evolution trials was generated from T4 bacteriophage in two steps. In the first step, we generated a new strain of T4, termed T4^pol_del^, in which the genes encoding the polymerase and the clamp loader complex are deleted. In the second stage, the genes encoding the RR2/T4 chimeric clamp-loader complex were introduced into T4^pol_del^ to generate the T4^chimera^.

#### Generation of T4^pol_del^

In our previous work, we developed a method to delete essential genes from the genome of bacteriophage T4. We had used the method to delete the genes encoding the clamp loader and the sliding clamp (Subramanian et al. 2021). Here, we apply the same method to generate T4^pol_del^, the strain of T4 in which the genes for the DNA polymerase and the clamp loader complex are deleted. The genes to be deleted are adjacent to each other on a ∼6 kb stretch of the T4 genome (32,886 bp – 30,341 bp, numbered according to the GenBank file with accession number AF158101). We used a CRISPR system to generate a cut in this span, and used homologous recombination to replace the span with a short, 20 bp segment to generate T4^pol_del^. The newly inserted 20 bp segment can be targeted using CRISPR, enabling us to insert the chimera at this site in order to generate _T4chimera._

We infected *E. coli* BL21 cells carrying three plasmids with wildtype T4 bacteriophage particles to generate T4^pol_del^. The first plasmid, termed the CRISPR T4 plasmid, encodes CRISPR-Cas12a and a guide RNA targeting the region to be deleted. The second plasmid, termed the donor plasmid, contains a new CRISPR-Cas12a recognizable site (TTTACCGGGAGGAAGATATAGCAC, PAM sequence is underlined) that is flanked on both sides by ∼1kb homologous arms. The homologous arms are identical to the regions on either side of the target locus on the T4 genome. The third plasmid, termed the helper plasmid, encodes a copy of the T4 genes that are to be deleted, and can therefore support the propagation of T4^pol_del^. We isolated T4^pol_del^ plaques and verified that the isolated phage is unable to propagate in bacteria unless the cells carry the helper plasmid. Sanger sequencing of the polymerase genetic locus in the isolated phage provided confirmatory evidence that we had indeed generated T4^pol_del^.

#### Generation of T4^chimera^

We constructed a new recombination plasmid, termed rec_loader plasmid, to introduce the chimera into T4^pol_del^. The rec_loader plasmid contains the gene encoding the T4 sliding clamp and the genes encoding the ATPase and clasp subunits of the chimera. These genes are flanked by arms of homology in order to recombine into the polymerase locus on the genome of T4^pol_del^ and generate T4^chimera^.

We infected *E. coli* BL21 cells carrying three plasmids with T4^pol_del^ to generate T4^chimera^. The first plasmid is the CRISPR plasmid that targets the newly inserted segment on T4^pol_del^. The second plasmid is the rec_loader plasmid discussed above that encodes the genes for the RR2/T4 chimeric clamp-loader complex. The third plasmid is the pol plasmid, a plasmid encoding the T4 DNA polymerase that is required to propagate T4^chimera^. We isolated phage particles the infection and used Sanger sequencing to verify the correct genomic integration of the genes corresponding to the chimera.

### Directed evolution through phage propagation

Directed evolution was performed by passaging phage populations in bacteria carrying a plasmid-borne copy of the D219A variant of the T4 DNA polymerase, an error-prone variant having a ∼1,000-fold higher mutation rate than the wildtype polymerase (Frey et al. 1993). Each trial of directed evolution was started with ∼10^5^ particles of T4^chimera^ added to 2ml of *E. coli* BL21 cells with the mutator plasmid (key resource table) that are exponentially growing with OD_600_ between 0.1 and 0.2. Phage was allowed to propagate in this culture for 8hrs to overnight at 37 ℃. The culture was centrifuged to precipitate cell debris and 1ml of the supernatant was saved for analysis. 2 µl of the supernatant was used to inoculate a fresh, 2 ml culture of bacteria with the mutator plasmid.

To perform Sanger sequencing, 1µl of the supernatant was used as template in a 20 µl PCR mix to amplify the clamp loader locus, using primers with sequences TAACCAAAATAGCGATTTTC and TTGCTGCTCAAATTGTTG.

### Molecular dynamics simulations

We modeled the AAA+ module of the RR2 clamp loader using Alphafold (Jumper et al. 2021). Molecular dynamics simulations were setup similar to our previous work (Subramanian et al. 2021). Histidine protonation states were inferred using H++ web server (Anandakrishnan et al. 2012). We solvated the model with TIP3P water within a truncated octahedron 15 Å larger than the AAA+ module and neutralized with Na+ using tleap (Case et al. 2019), and added 150mM NaCl based on this initial volume. Six independent trajectories with different random seeds were started with this model. In each case, 1ns Langevin dynamics (1/ps friction coefficient) with 1fs time step was used to equilibrate the system in NVT at 310.15K within pmemd on an NVIDIA A40 GPU. After this, a Monte Carlo barostat was used with 2fs time step to equilibrate the NPT for 1ns. Then, the models were subjected to equilibration for a total of 2 µs and distances were between the guanidino carbon of Arg 85 and the Cγ of Asp 86 were calculated for the equilibration trajectory.

## Acknowledgements

We thank members of the Kuriyan lab for helpful discussions and Kent Gorday for help in setting up the molecular simulations. We thank Prof. Jim Karam, Emeritus Professor, Tulane University for generously sharing RB69 and RR2 phages with us. WZ thanks the Chemistry Visiting Students program for funding support.

---

**Supplementary Figure 1.**
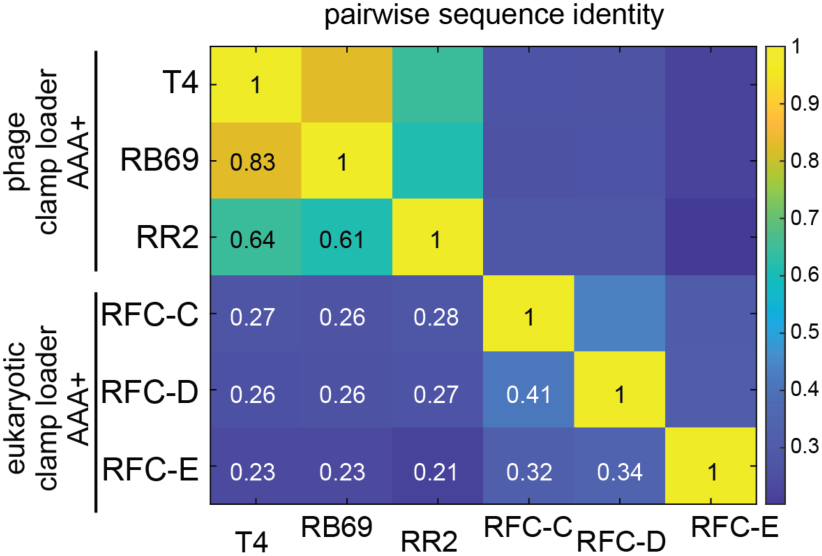
Sequence similarity between AAA+ modules in the constructed chimeric clamp loaders. Pairwise sequence identitity between AAA+ modules used in the study.

**Supplementary Figure 2.**
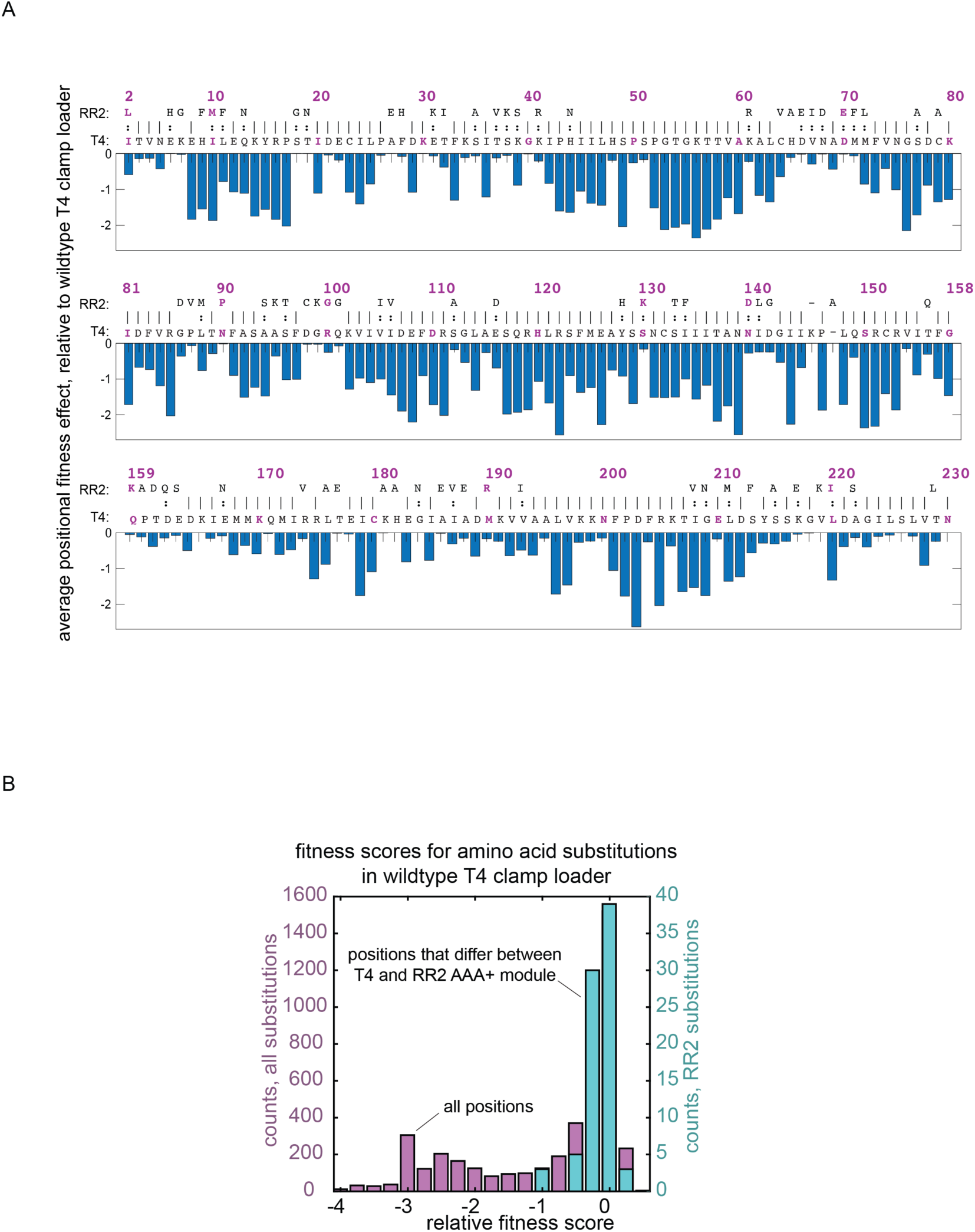
Mutational effects in the AAA+ module of the wildtype T4 clamp loader. (A) Average effect mutating each position of the AAA+ module in the wildytpe T4 clamp loader, based on fitness values from (Subramanian et al. 2021)).The sequence of the RR2 AAA+ module is aligned with T4 AAA+ module. (B) Distribution of fitness effects, for the wildtype T4 clamp loader, from introducing individual substitions to each of the differing residues in the corresponding RR2 AAA+ module.

**Supplementary Figure 3.**
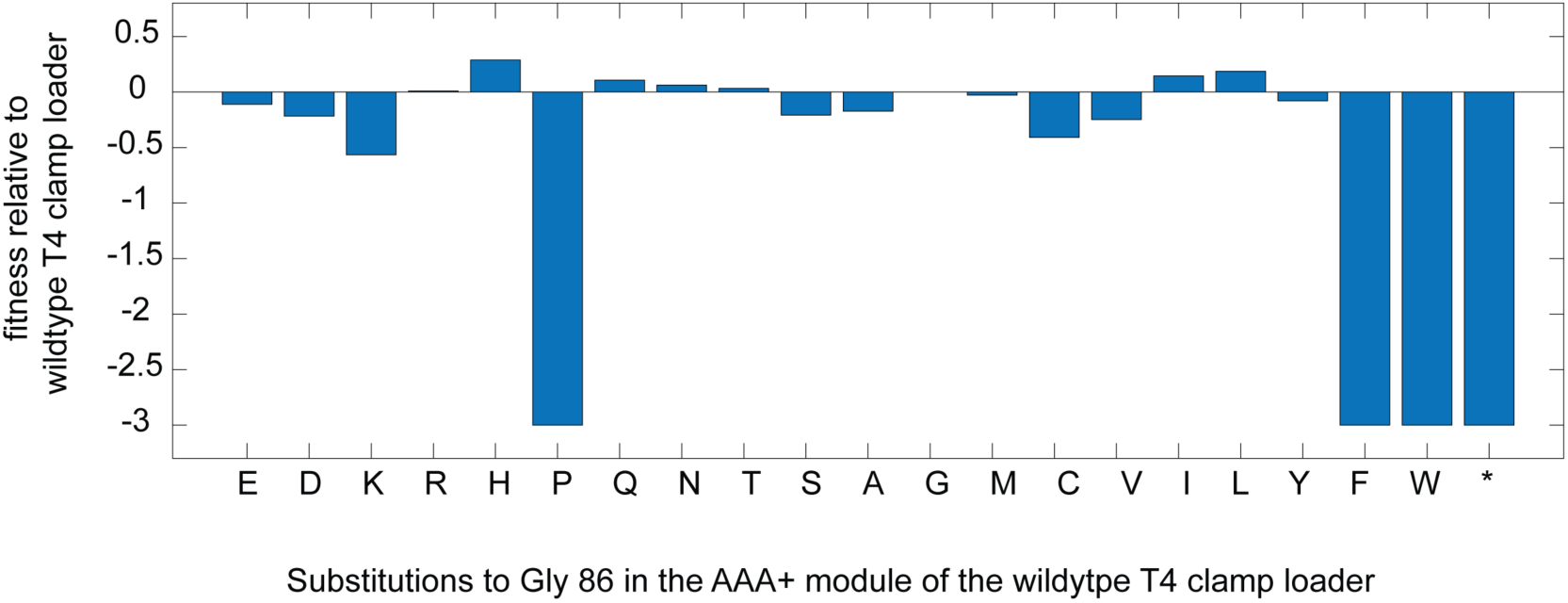
Effect of single amino acid substitutions at position 86 of the ATPase subunit of the wildtype T4 clamp loader. Fitness values from (Subramanian et al. 2021)).

**Supplementary Figure 4.**
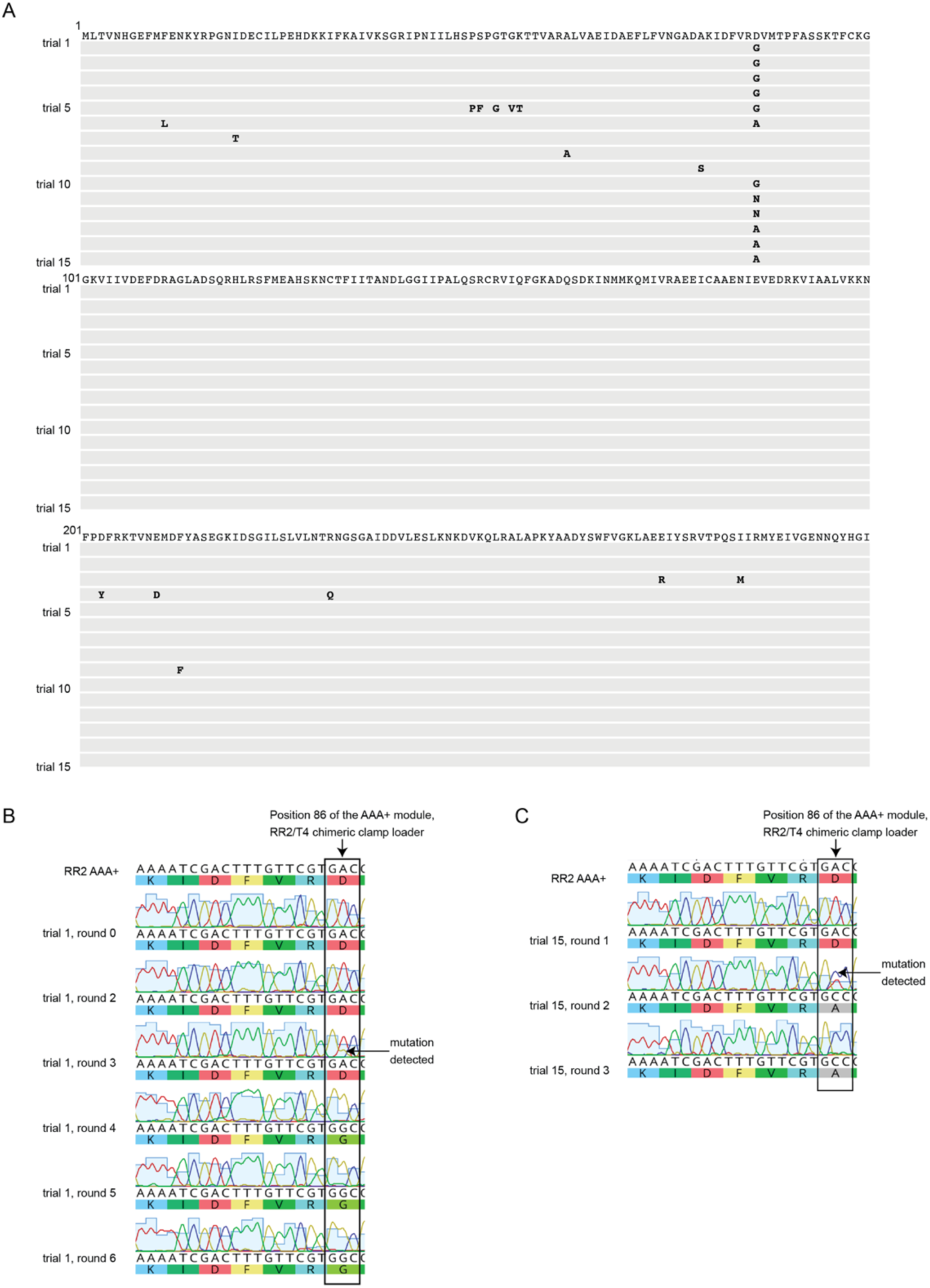
Sanger Sequencing of *in vitro* evolution trials. (A) Sequencing of the AAA+ module of the RR2/T4 chimeric clamp loader locus in T4^chimera^, after round 10 from each of the 15 parallel trials of *in vitro* evolution. Changes to the starting protein sequence is shown. (B) and (C) Sanger sequencing chromatograms for the early rounds of trial 1 and trial 15, for the 21-nucleotide segment ending at the codon corresponding to residue 86 of the AAA+ module of the RR2/T4 chimeric clamp loader locus.

**Supplementary Figure 5.**
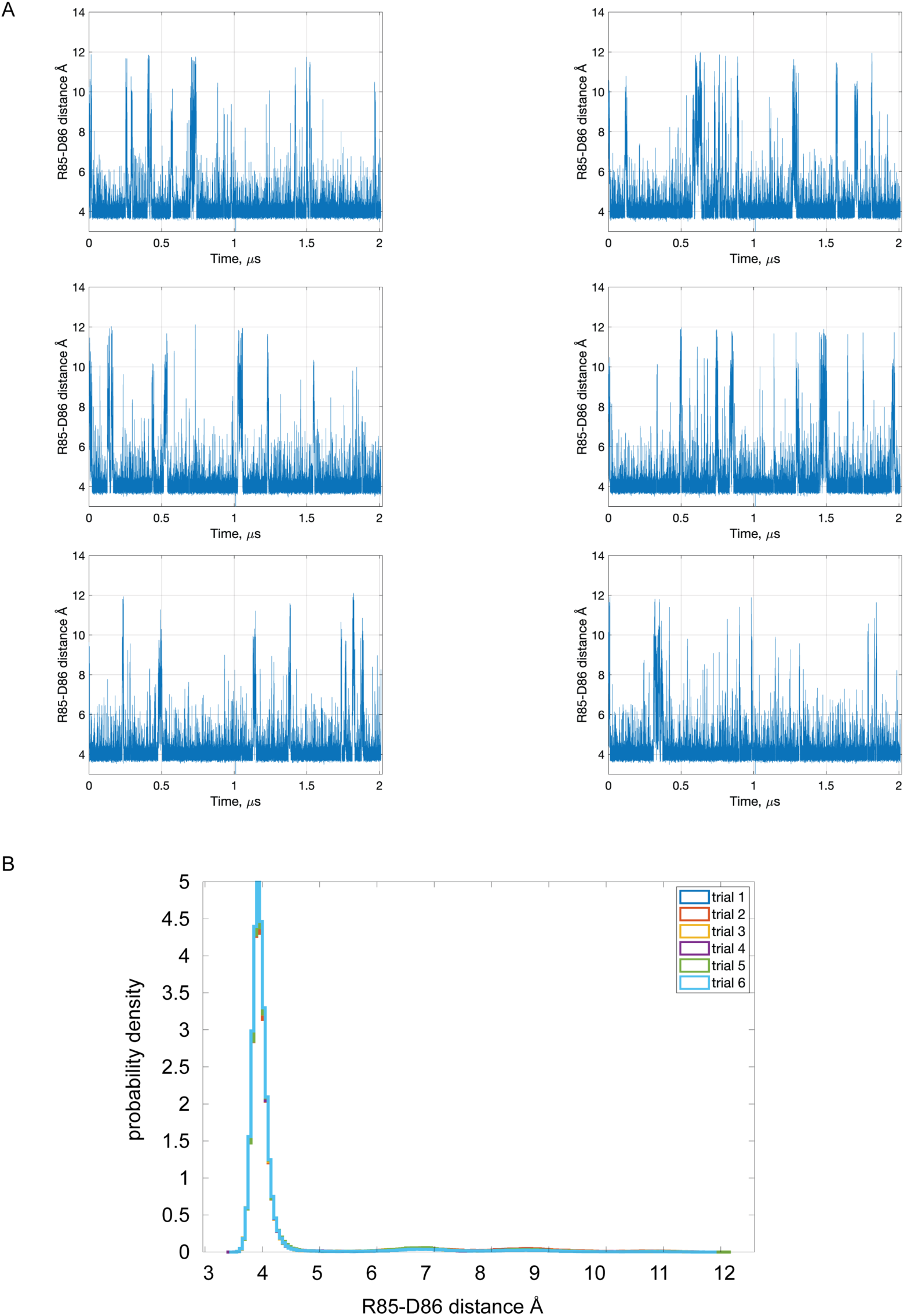
Molecular dynamics simulations of the AAA+ module of the RR2 clamp loader. (A) The distance between the guanidino group of Arg 85 and the γ-carbon (sidechain carboxyl group) of Asp 86 are plotted for a 2µs simulation for six molecular dynamics trajectories. (B) Probability density of the distance between Arg 85 and Asp 86, from **A**.

**Supplementary Figure 6.**
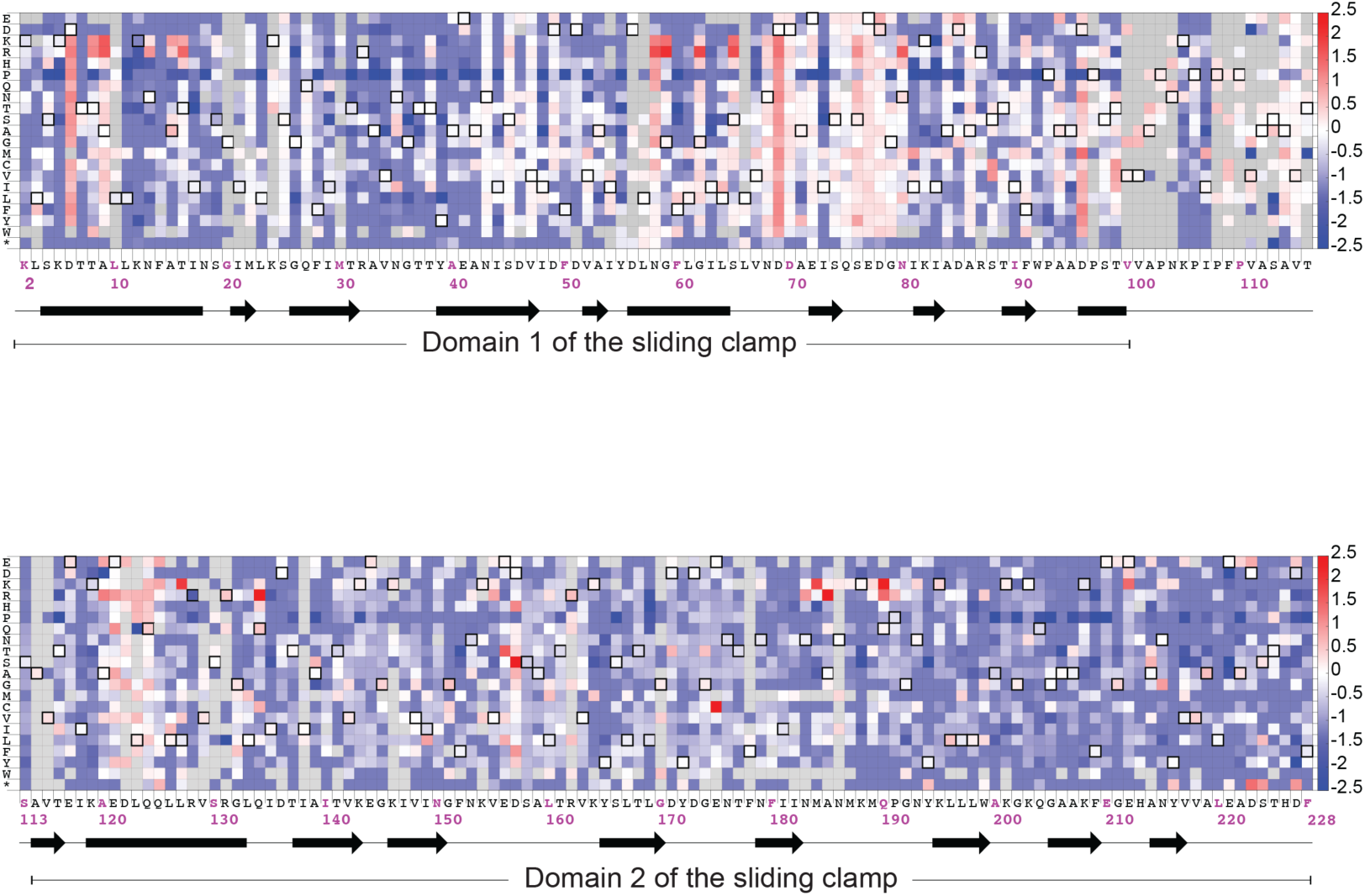
Mutational sensitivity of the T4 sliding clamp in the context of the RR2/T4 chimeric clamp loader. Fitness scores are calculated relative to the unmutated T4 sliding clamp in the context of the RR2/T4 chimeric clamp loader. Amino acid substitutions that result in the wild-type sequence are denoted by pixels with a black border. Gray pixels denote mutations with insufficient counts (fewer than 10 counts) when sequencing the plasmid library. The secondary structure of the wild-type sequence is indicated below the wild-type sequence, with α helices as rectangles and β strands as arrows.

**Supplementary Figure 7.**
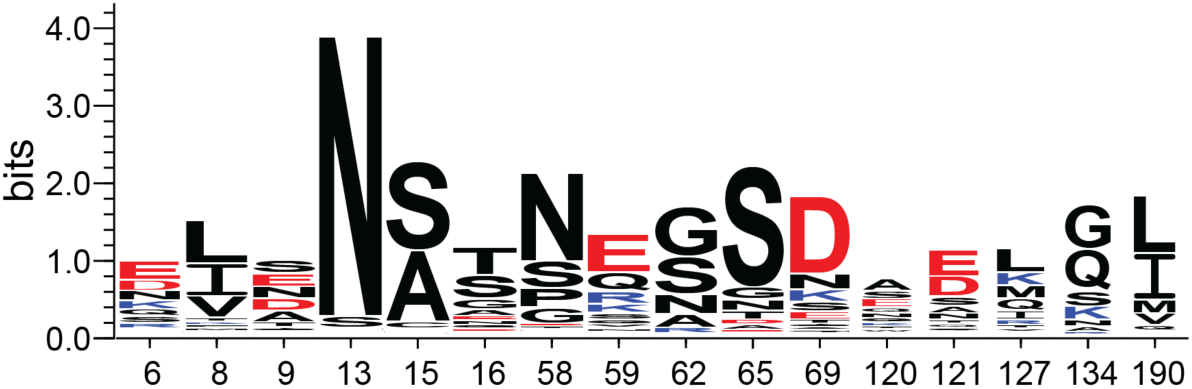
Sequence logo of the 16 positions in the T4 sliding clamp in which substitutions can result in at least a 10-fold increase in phage propagation. Residues with acidic side chains (Asp and Glu) are shown in red, residues with basic sidechains (Arg and Lys) are shown in blue. Asn 13, a residue in domain 1 of the sliding clamp that forms a hydrogen-bonded interaction with the amide backbone of domain 2 at residue 204 in the T4 clamp loader, shows a sequence signature of strong evolutionary conservation.

**Supplementary Figure 8.**
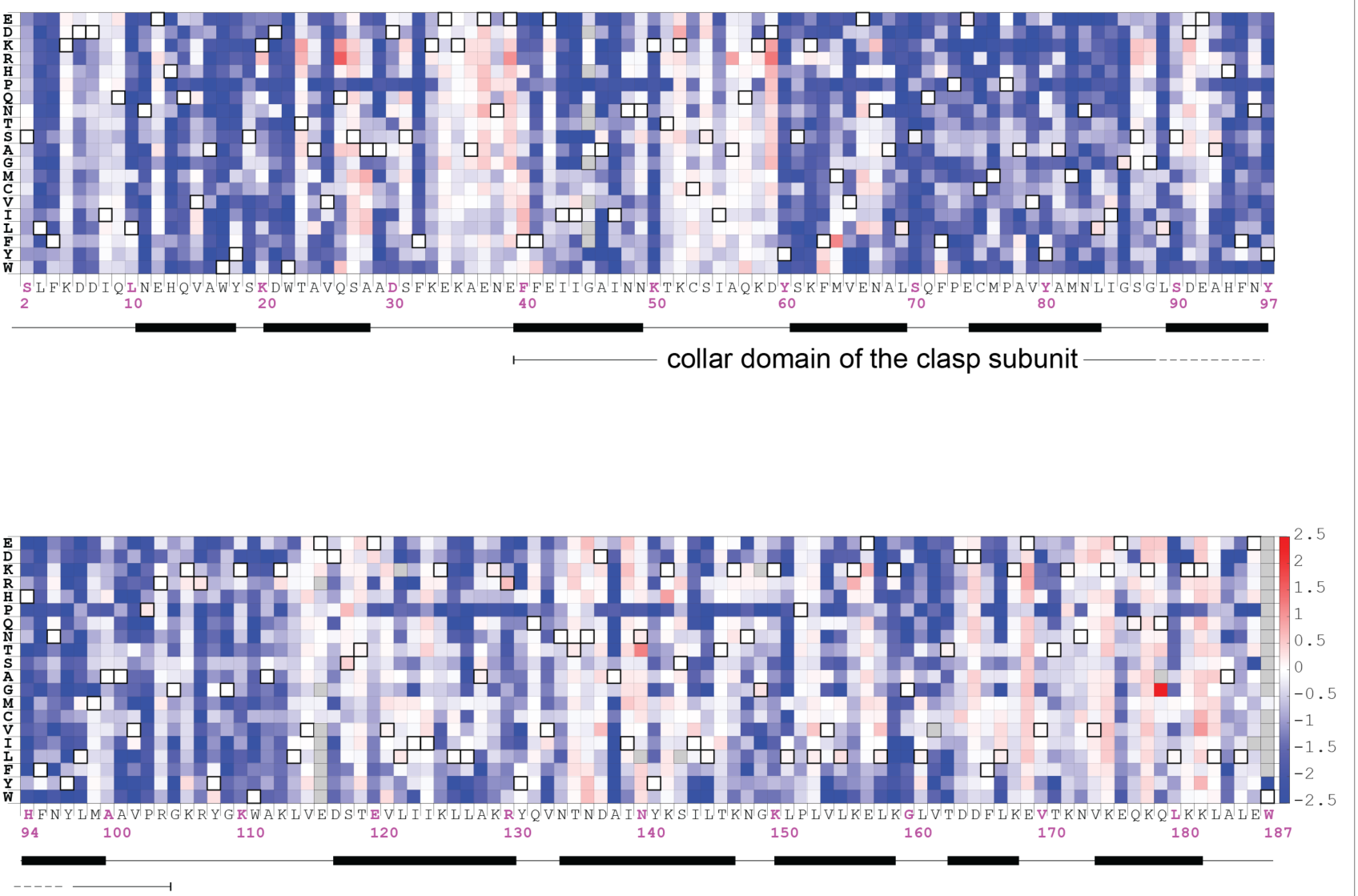
Mutational sensitivity of the T4 clasp subunit in the context of the RR2/T4 chimeric clamp loader. Fitness scores are calculated relative to the unmutated RR2/T4 chimeric clamp loader. Amino acid substitutions that result in the wild-type sequence are denoted by pixels with a black border. The secondary structure of the wild-type sequence is indicated below the wild-type sequence, with α helices as rectangles and β strands as arrows.

**Supplementary Figure 9.**
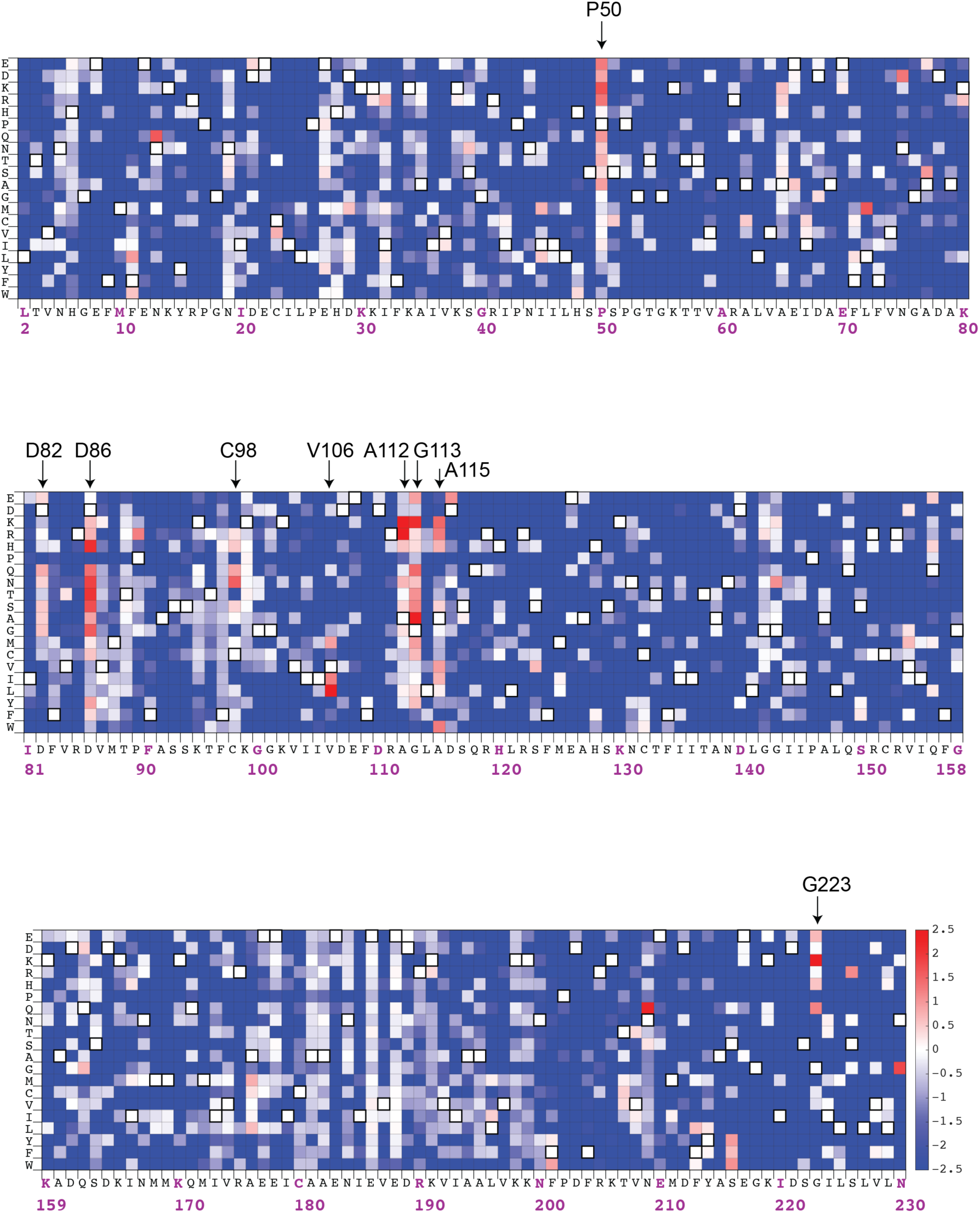
Mutational sensitivity of the AAA+ module of the RR2/T4 chimeric clamp loader. Fitness scores are calculated relative to the unmutated RR2/T4 chimeric clamp loader. Amino acid substitutions that result in the wild-type sequence are denoted by pixels with a black border. The secondary structure of the wild-type sequence is indicated below the wild-type sequence, with α helices as rectangles and β strands as arrows.

**Supplementary Figure 10.**
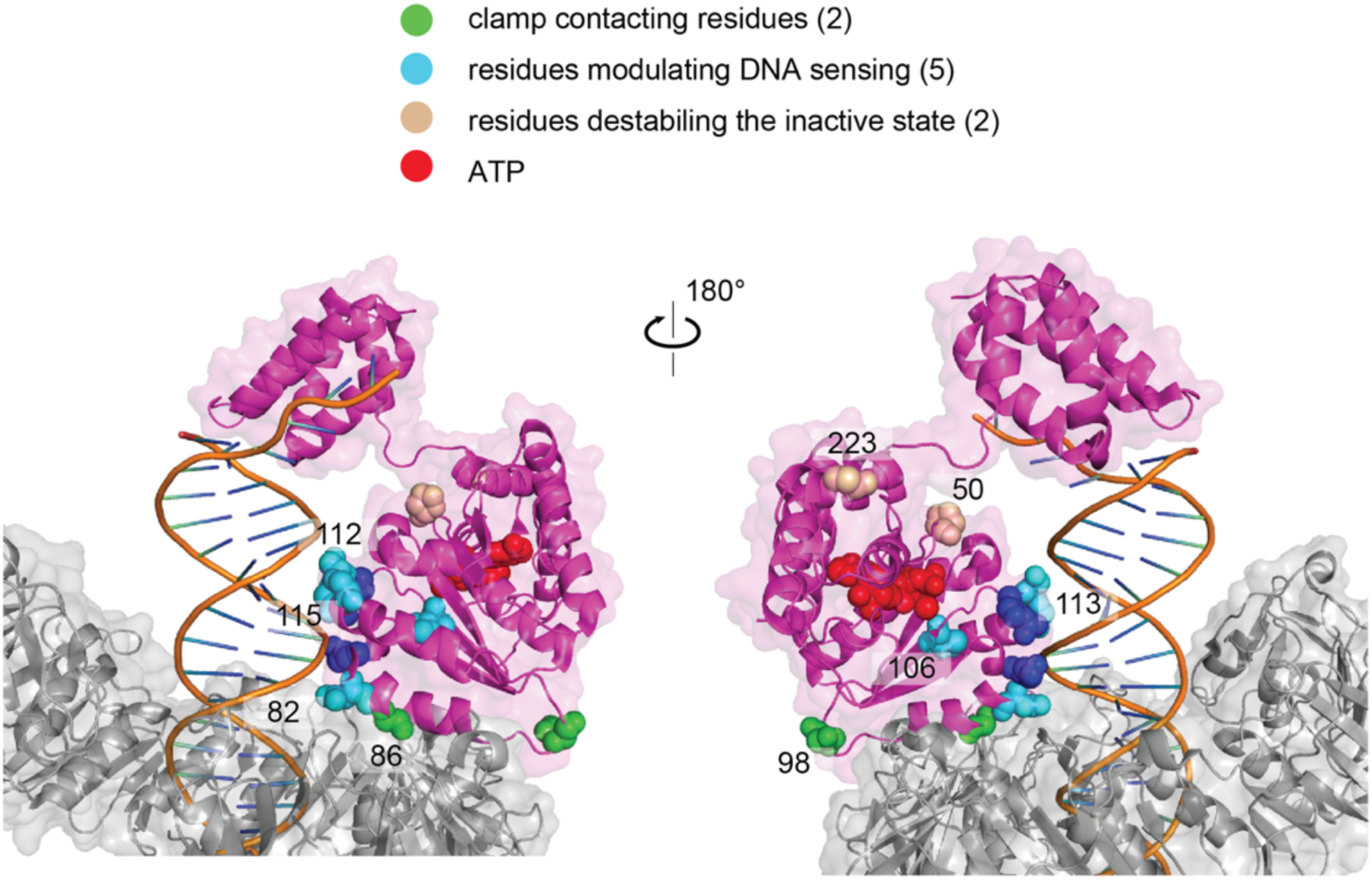
Gain-of-function mutations in the AAA+ module of the RR2/T4 chimeric clamp loader. The spatial location of residues in the AAA+ module of the RR2/T4 clamp loader in which substitutions can result in at least 10-fold increase in phage propagation (PDB ID: 3U60). Residues are grouped into 3 set: clamp contacting residues, DNA contacting residues and residues that can destabilize the DNA-free inactive state of the clamp loader.

**Supplementary Figure 11.**
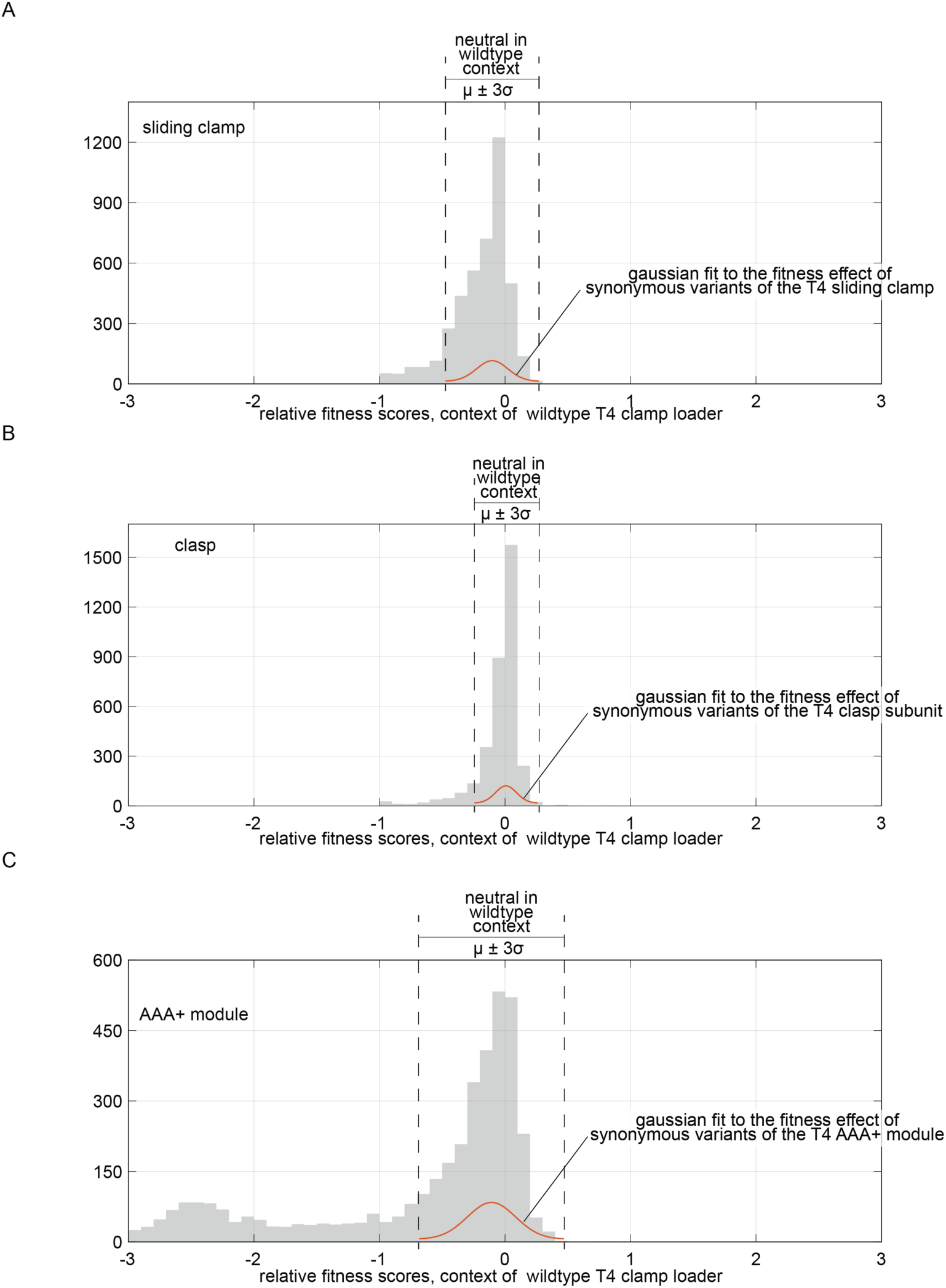
Determining the threshold for neutral mutations in the wildytpe T4 clamp-loader complex. The distribution of fitness effects of point mutations to the sliding clamp (A), the clasp (B) and the AAA_ module (C) of the wildytpe T4 clamp-loader complex are plotted as histograms (gray). A gaussian fit to the fitness effect of synonymous variants of the reference sequence is shown in red, along with the three standard deviations threshold for the fit.

